# Within-host antigenic selection of influenza A virus dominates over stochasticity but is limited by fitness tradeoffs and timing of the immune response

**DOI:** 10.64898/2026.08.19.745600

**Authors:** Vedhika Raghunathan, Christina M. Leyson, Matthew Gaddy, Lucia Ortiz, Nahara Vargas-Maldonado, Jens Wrammert, Georgii A. Bazykin, Daniel B. Weissman, David VanInsberghe, Anice C. Lowen

**Affiliations:** Department of Microbiology and Immunology, Emory University School of Medicine, Atlanta, GA, USA; Division of Infectious Diseases, Department of Pediatrics, Emory University School of Medicine, Atlanta, GA, USA; Department of Biology, Emory University, Atlanta, GA, USA; Department of Physics, Emory University, Atlanta, GA, USA

## Abstract

Despite antigenic evolution at the global scale, positive selection of influenza virus antigenic variants is not readily observed within hosts. Here, we tested the extent to which fitness tradeoffs, the timing of immune pressure, and stochastic effects impede antigenic selection within pre-immune hosts. We used genetically barcoded influenza A/Texas/50/2012 (H3N2) viruses (Tx/12) in a guinea pig model to probe these dynamics. Positive selection of an antigenic variant was reliant on a high strength of immune pressure acting early in infection. However, when fitness tradeoffs of the antigenic change were lessened, a lower strength and later introduction of immune pressure favored the antigenic variant. In all conditions, barcode dynamics revealed moderate stochastic effects. Our results suggest that stochastic evolution does not impede selection during acute influenza virus infection. The rarity of antigenic escape may instead stem from low mutational supply, fitness tradeoffs, and the intrinsic delay between infection and antibody recall.

## INTRODUCTION

The evolution of influenza A virus (IAV) leads to the global circulation of antigenically novel variants every few years. Antibodies generated in response to infection or vaccination primarily target the two surface glycoproteins of IAV, hemagglutinin (HA) and neuraminidase (NA) [1–5]. These antibodies drive the emergence of antigenic variants that escape humoral responses and therefore outcompete ancestral viruses [1–6]. The result is a clear pattern of recurring selective sweeps at the scale of the global human population [1–6].

Each antigenic variant that spreads globally arises initially within an infected host. Nevertheless, the positive selection of antigenic variants within acutely infected humans has not been readily observed. Studies of naturally infected individuals have shown that influenza virus populations within hosts are typically low in diversity, with fewer than 10 variants present at greater than 2% frequency [7]. Substitutions in antigenic sites furthermore remain at low frequencies over the course of infection and are not more commonly found than substitutions in non-antigenic sites [7–10]. While evidence of positive selection is rare, low rates of non-synonymous substitutions suggests that purifying selection is active [7–11] and a prominent role for stochastic effects has been noted [7–10]. In particular, estimates of effective population size (N_e_) [12] in the range of 50-300 point to an important role for genetic drift in driving changes in population composition over time [13, 14]. While evolutionary dynamics can be distinct at different spatial scales, these observations at a within-host level draw a sharp contrast with patterns observed at the host population level.

Given the efficiency of purifying selection within hosts, the fitness costs often associated with antigenic variants may impede their positive selection. In HA, mutations that enable antibody escape often also modulate viral binding to cell surface receptors and can be deleterious [15, 16]. While this effect can be alleviated through viral acquisition of epistatic mutations [15, 17–20], HA function constrains its antigenic evolution.

Stochastic evolutionary change can also impede selection. Selection will only be observed where its deterministic effects outweigh the magnitude of stochastic fluctuations in allele frequencies. The relative timescales of these two processes are also important: for example, if an antigenic variant is lost early through genetic drift [7, 21, 22], memory immune responses that are recalled in the days after infection [23, 24] may act too late to impose selection.

Finally, limited viral diversity may account for the rarity of positive selection at the within-host level. Transmission of IAVs reduces viral diversity, such that the diversity that establishes in a new host is low, irrespective of the diversity that was present in the donor host. This loss of diversity has been attributed to either stochastic bottlenecks [7, 21, 22, 25] or selection [10, 26–29] acting during transmission. Through either mechanism, the resultant paucity of variants within the host limits the potential for antigenic evolution. Indeed, the positive selection of IAV variants within hosts has been noted in the context of experimental inoculations, which are not typically characterized by stringent bottlenecks during the establishment of infection [26, 30, 31].

With the application of a well-controlled experimental model, here we sought to define the conditions that prevent or enable positive selection of IAV antigenic variants within hosts. We used influenza A/Texas/50/2012 (H3N2) virus and two antigenic variants thereof, one which incurred a fitness cost and one in which that cost was offset by two compensatory mutations. Each virus also contained a neutral barcode to monitor stochastic effects. Working in a pre-immune guinea pig model, we found that selection was observed consistently but favored the antigenic variant only when the fitness tradeoff was overcome by strong, early immune pressure or when the tradeoff was eased through second-site mutations. In contrast, stochastic effects were moderate across all conditions such that neutral variation was often maintained.

## RESULTS

### Design and characterization of barcoded viruses

To monitor antigenic selection within hosts, we used two variants of influenza A/Texas/50/2012 (H3N2) virus (Tx/12): the wild type (WT) and HA F159S, a previously characterized antigenic variant [16]. To monitor stochastic effects that could also shape viral populations, each virus was outfitted with a distinct genetic barcode. In this way, each antigenic variant was represented by thousands of unique genotypes. Barcodes were placed within the HA gene and comprised 10 synonymous polymorphisms that occurred naturally among human influenza A/H3N2 viruses circulating from 2003-2020. This design was chosen to avoid viral attenuation and fitness differences between barcode genotypes [21, 32]. With 10 bi-allelic sites within a 110-nucleotide region of the HA segment, each library comprised 2^10^ = 1,024 unique barcodes (Figure 1A). We named the viruses modified in this manner Tx/12 HA WT BC1 and Tx/12 HA F159S BC2.

**Figure 1:**
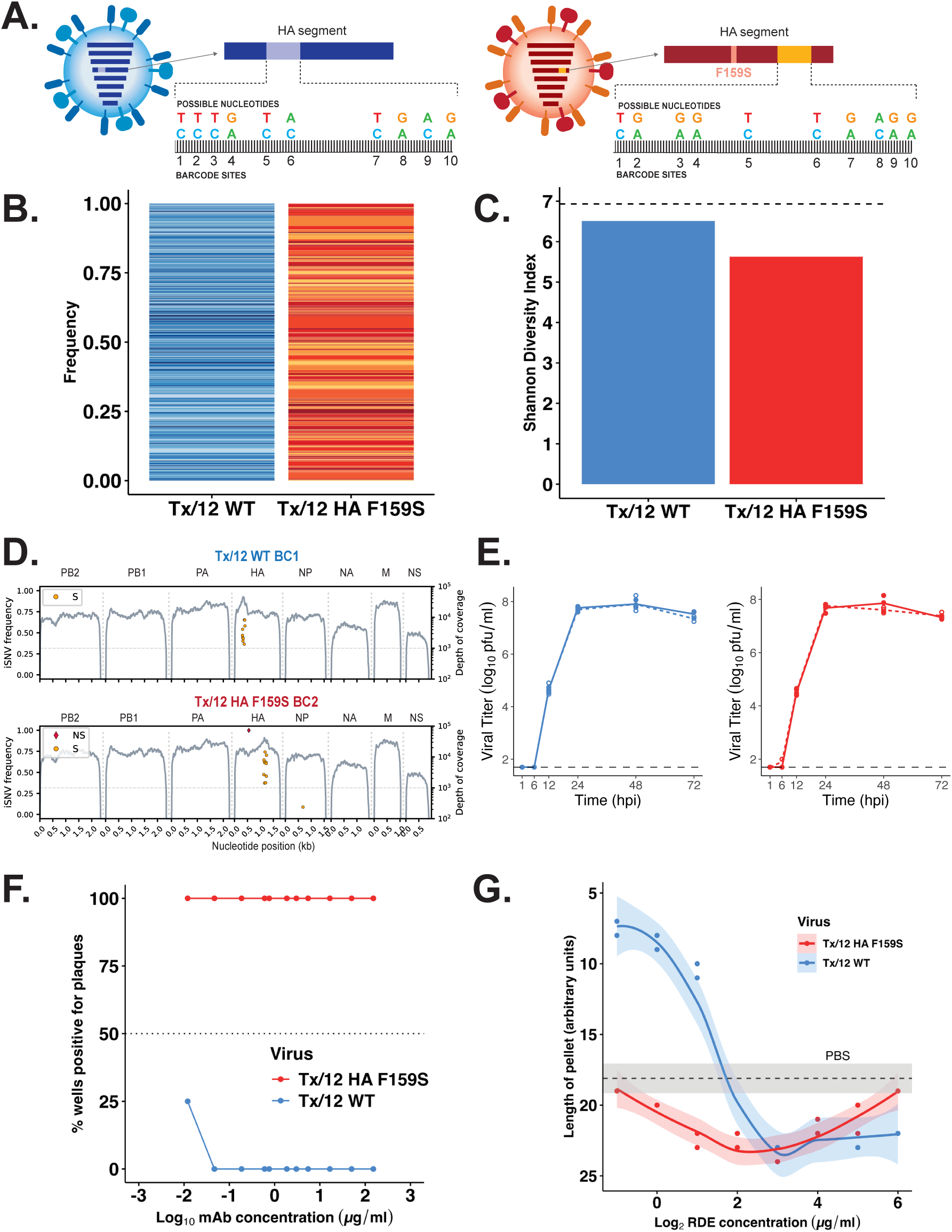
Tx/12 HA WT and HA F159S barcoded viruses are tools to monitor selective and stochastic effects within hosts. **(A)** Barcode design of HA segments of influenza A/Texas/50/2012 (H3N2) viruses (Tx/12). Tx/12 WT and HA F159S each contain a distinct barcode region in the HA gene, within which there are 10 bi-allelic sites. **(B)** Barcode frequencies of passage 1 virus stocks diluted 1:10. Each colored section represents a different barcode, and its height is the relative frequency within the sample. **(C)** Shannon diversity of barcoded passage 1 virus stocks diluted 1:10. **(D)** Whole genome sequencing of barcoded virus stocks. Categorized iSNVs (S=synonymous, NS=nonsynonymous) are plotted at their frequency on the left axis and sequencing coverage (gray line) is plotted on the right axis. Segments are identified above each trace. **(E)** Multicycle growth curves of Tx/12 HA WT (blue, solid line), Tx/12 HA WT BC1 (blue, dashed line), Tx/12 HA F159S (red, solid line), and Tx/12 HA F159S BC2 (red, dashed line). Infections were performed in triplicate, shown as individual points with the line plotted through the mean. Dashed line represents limit of detection of 50 pfu/ml. Repeated measures ANOVA (p=0.088 and 0.276 for WT and HA F159S, respectively) was performed. **(F)** Plaque reduction neutralization of Tx/12 HA WT and Tx/12 HA F159S using mAb 2A05. Data are expressed as the percent of wells within each quadruplicate that were positive for ≥1 pfu. Dashed line represents 50% (2/4 replicates) having ≥1 pfu. **(G)** Binding of Tx/12 WT and HA F159S viruses to receptor destroying enzyme (RDE)-treated red blood cells. Data are expressed as length of red blood cell pellet (performed in duplicate). Solid lines are polynomial regression estimates fit to the data with shaded areas showing 95% confidence intervals. Dashed line represents mean pellet length in PBS negative control with shaded 95% confidence interval.

The standing barcode diversity of our passage 1 (P1) viral stocks was evaluated by next generation sequencing. All 1,024 barcodes were detected for Tx/12 HA WT BC1 while 998 were detected for Tx/12 HA F159S BC2 (Figure 1B). Shannon diversity was 6.5 and 5.6 for Tx/12 HA WT BC1 and Tx/12 HA F159S BC2, respectively, with the reduction relative to the theoretical maximum value of 6.93 attributable to minor variation in the relative abundance of constituent barcodes (Figure 1C). Whole genome sequencing of these P1 stocks revealed an absence of *de novo* mutations that would be expected to alter barcode dynamics (Figure 1D) and analysis of multicycle replication in MDCK cells showed no impact of the barcodes (Figure 1E).

HA F159S has been shown to mediate escape from neutralization by mAb 2A05 [33]. To confirm this phenotype, plaque reduction neutralization assays were performed with mAb 2A05 and Tx/12 WT and HA F159S viruses. While the WT virus was neutralized, Tx/12 HA F159S showed strong escape (Figure 1F). The HA F159S change has also been shown to incur a fitness cost by lowering cell binding avidity [16]. A cell binding avidity assay confirmed this pleiotropic effect: compared to the WT virus, the HA F159S variant bound poorly to red blood cells (Figure 1G).

### Positive selection of Tx/12 HA F159S is reliant on strong immune pressure

To evaluate the efficiency of within-host selection under varying strengths of immune pressure, animals were co-infected with Tx/12 HA WT BC1 and Tx/12 HA F159S BC2 viruses. Initial subpopulation frequencies were either ∼10:1 or ∼1:1 (Supplemental Table 1). Immune pressure was modeled by instilling mAb 2A05 into the nasal cavity daily [34], and the strength of pressure was varied by using a range of mAb doses. To monitor changes in viral population composition, daily nasal lavage samples were collected. Antigenic variant abundance in these samples was evaluated by quantifying HA genes encoding F or S at position 159, while barcode composition within each subpopulation was assessed using next generation sequencing.

Results were similar with 10:1 and 1:1 starting populations (shown in Figure 2 and Supplemental Figure 1, respectively). At no or low doses of mAb 2A05, Tx/12 HA F159S BC2 was consistently lost during infection. At intermediate doses of mAb 2A05, the fate of Tx/12 HA F159S BC2 varied between animals: it was either lost, as seen with low doses of mAb, or it underwent an initial increase in frequency followed by a decline. Finally, at high doses of mAb 2A05, Tx/12 HA F159S BC2 was rapidly fixed (Figure 2A and Supplemental Figure 1A). To quantify these trends, the growth rate of Tx/12 HA F159S BC2 was estimated. The growth rate (*β*) was defined as the change in 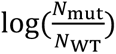 per day*N*_WT_ post-infection, where *N*_mut_ and *N*_WT_ are the barcode read counts from the Tx/12 HA F159S BC2 and Tx/12 HA BC1 viruses, respectively. Positive values of *β* indicate an increase in the frequency of Tx/12 HA F159S BC2 whereas negative values indicate a decrease. At no or low doses of mAb 2A05, *β* was <0. At intermediate doses of mAb 2A05, *β* estimates approached 0, but only at high doses of mAb 2A05 was *β* >0 (Supplemental Figure 2A). Thus, increased immune pressure was essential to enable efficient antigenic selection.

**Figure 2:**
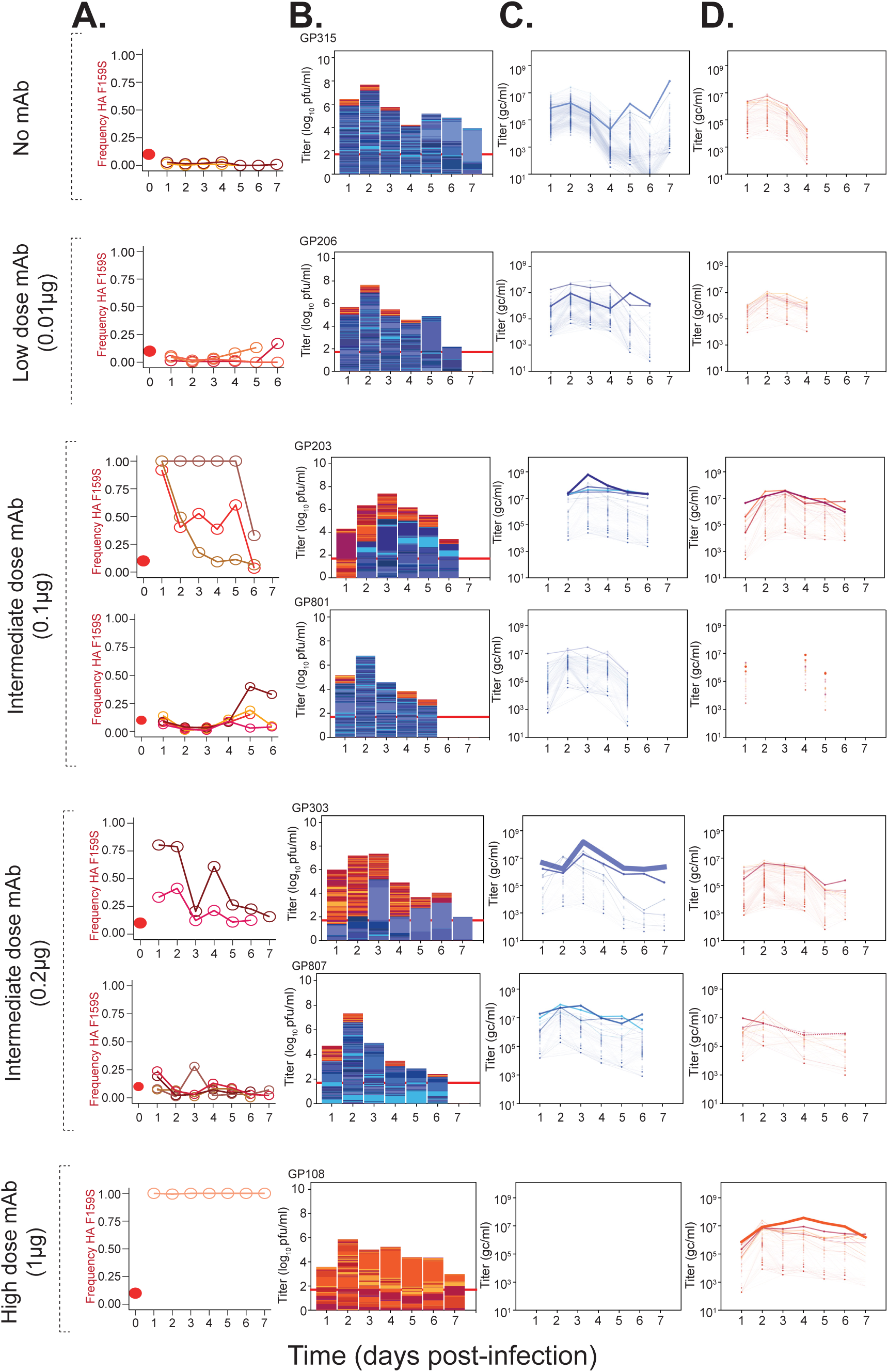
Deterministic forces shape overall antigenic variant abundance despite some concurrent stochasticity within subpopulations. Tx/12 HA WT BC1 virus is represented by blue/purple colors and Tx/12 HA F159S BC2 is represented by red/yellow colors. Animals inoculated with 90% WT and 10% HA F159S BC viruses. The mAb dose applied daily is indicated at the left of each row. **(A)** Frequency of the Tx/12 HA F159S allele over days of infection. Solid dot denotes starting frequency of Tx/12 HA F159S in the inoculum. Each shade of red represents a different animal. Animals in which *de novo* mutations were associated with barcode dynamics are not displayed. Panels **(B)-(D)** contain a representative animal per group, with the same animal featured in all panels. Data from all animals are shown in Supplemental Figure 3. **(B)** Stacked bar plots where the total height of the column is scaled to viral titer (pfu/ml). Each colored section within the column represents a unique barcode, and its height is the relative frequency within the sample. The total height of red/yellow bars and blue/purple bars is scaled to the genome copies/ml (gc/ml) titer of each virus. Red horizontal line represents limit of detection of plaque assay (50 pfu/ml). **(C)** and **(D)** show Tx/12 HA WT and F159S individual barcodes over time respectively. Each line is a unique barcode with its frequency scaled to the gc/ml titer. Line thickness is scaled to the average frequency of the barcode. Gaps denote results below the limit of detection. Dashed lines connect data separated by single days below the limit of detection.

Under all conditions, two patterns of barcode dynamics were apparent (Figure 2B-D; Supplemental Figure 1B-D; Supplemental Figure 3). In some animals, many barcodes within the predominant antigenic subpopulation were maintained, showing that stochastic effects were muted. Conversely, in other animals, the dominance of a particular antigenic subpopulation was driven by a few barcode lineages, with the remaining sister lineages declining in frequency over time. Importantly, however, even in this latter group where stochastic effects were apparent, this stochasticity did not preclude selection from acting.

### Delayed immune pressure restricts the efficiency of antigenic selection

Even in individuals with pre-existing immune memory, the recall of immune effectors takes time. We therefore evaluated the extent to which the observed selective processes depended on the timing of immune pressure. Animals were again co-infected with a mixture of Tx/12 HA WT BC1 and Tx/12 HA F159S BC2 viruses at ratios of ∼10:1 or ∼1:1 (see Supplemental Table 1 for inoculum compositions). A high dose of mAb 2A05 sufficient to favor Tx/12 HA F159S BC2 when delivered at the time of inoculation was instilled intranasally at 0 h, 6 h, 24 h, or 48 h post-inoculation. Following this initial delay, all animals received the same high dose of mAb 2A05 daily. As before, nasal lavage samples were collected daily and analyzed to define antigenic variant abundance and barcode composition.

When there was no delay between inoculation and initial instillation of mAb 2A05, Tx/12 HA F159S BC2 was quickly fixed (Figure 3A and Supplemental Figure 4A). Correspondingly, *β* was > 0 under this condition (Supplemental Figure 2B). When immune selection was initiated 6 h, 24 h, or 48 h after inoculation, one of two trends was observed: either the frequency of Tx/12 HA F159S BC2 increased following a delay, or it dropped such that this variant was either lost or remained a minor constituent throughout infection (Figures 3A and Supplemental Figure 4A). With delayed mAb delivery, *β* estimates were significantly reduced and not significantly different from zero (Supplemental Figure 2B). Thus, immune pressure was more effective in enabling positive antigenic selection during the establishment of infection than during subsequent rounds of viral replication.

**Figure 3:**
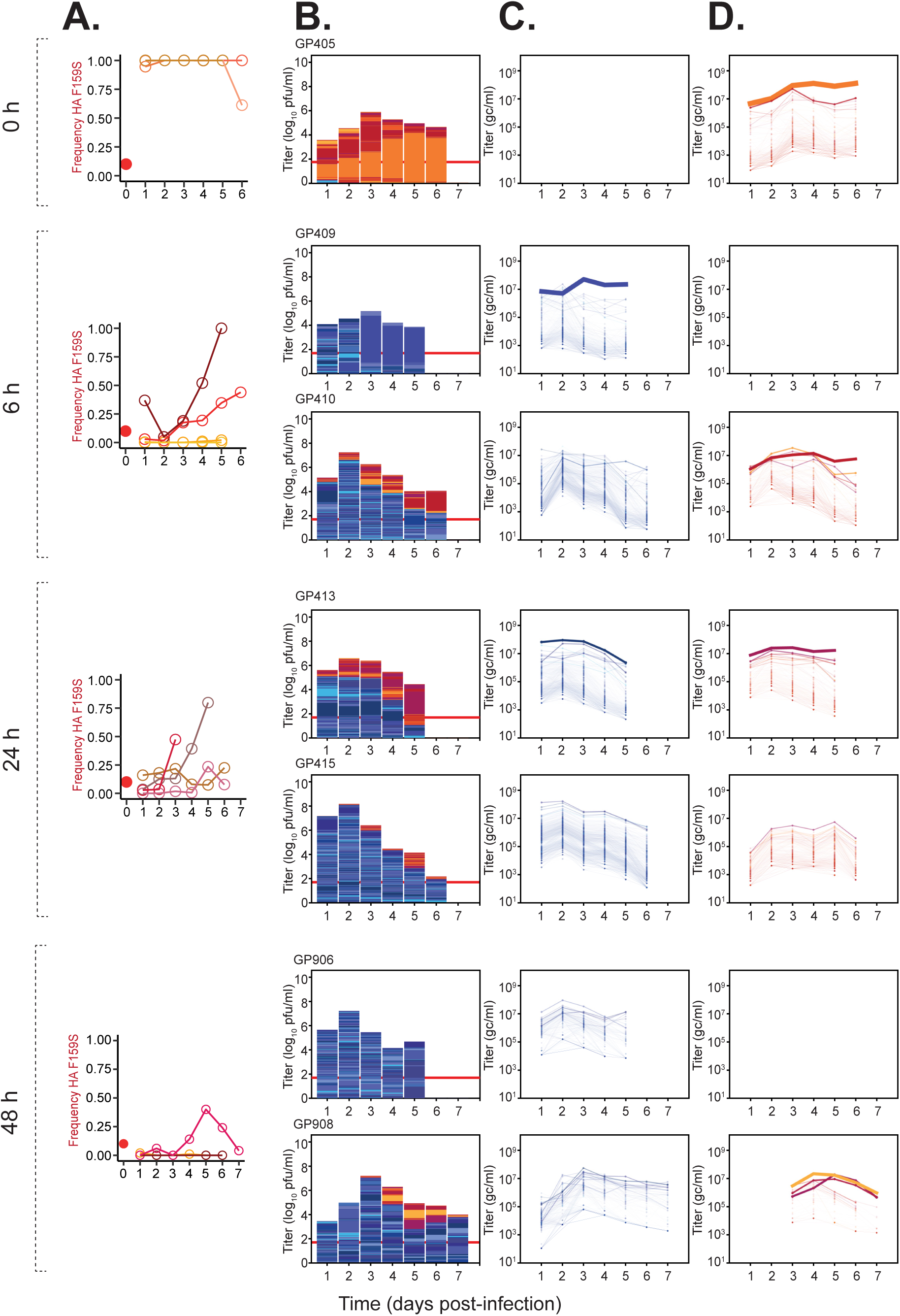
Delay of mAb treatment leads to delayed and inconsistent antigenic selection. Tx/12 HA WT BC1 virus is represented by blue/purple colors and Tx/12 HA F159S BC2 is represented by red/yellow colors. Animals were inoculated with 90% WT and 10% HA F159S BC viruses. The time interval in hours between viral inoculation and the first administration of a high dose of mAb (0.4 ug) is indicated at the left of each row. **(A)** Frequency of the Tx/12 HA F159S allele over days of infection. Solid dot denotes starting frequency of Tx/12 HA F159S in the inoculum. Each shade of red represents a different animal. Animals in which *de novo* mutations were associated with barcode dyanmics are not displayed. Panels **(B)-(D)** contain a representative animal or animals per group. The same animal is featured in all panels. Two representative animals are shown where a group was characterized by two distinct outcomes. Data from all animals are shown in Supplemental Figure 5. **(B)** Stacked bar plots where the total height of the column is scaled to viral titer (pfu/ml). Each colored section within the column represents a unique barcode, and its height is the relative frequency within the sample. The total height of red/yellow bars and blue/purple bars is scaled to the gc/ml titer of each virus. Red horizontal line represents limit of detection of plaque assay (50 pfu/ml). **(C)** and **(D)** show Tx/12 HA WT and F159S individual barcodes over time respectively. Each line is a unique barcode with its frequency scaled to the gc/ml titer. Line thickness is scaled to the average frequency of the barcode. Gaps denote results below the limit of detection.

The trends in barcode dynamics observed in these animals mirrored those seen when selection was imposed at the time of inoculation: in some animals, many barcodes within the dominant antigenic subpopulation were maintained, while in others, the positive selection of either virus was driven by a few component lineages, revealing concurrent stochastic effects (Figure 3B-D; Supplemental Figure 4B-D; Supplemental Figure 5). As before, however, even when stochastic effects were prominent, they did not prevent selection from defining the relative subpopulation frequencies within the host.

### *De novo* mutations near the receptor binding site arise in Tx/12 HA F159S BC2

To determine if barcode dynamics were linked to *de novo* mutations, viral whole genome sequencing was performed on a subset of nasal lavage samples. A total of 72 guinea pigs across the different experimental conditions were chosen, and three to four nasal wash samples spanning the infection period were sequenced from each. All animals that showed any barcodes reaching ≥ 10% frequency were included in this analysis and one animal from each experiment that maintained high barcode diversity was included for comparison. As an additional comparator, samples derived from groups of four guinea pigs that had been mono-infected with Tx/12 HA WT BC1 or Tx/12 HA F159S BC2 were subjected to viral whole genome sequencing.

In all guinea pigs, *de novo* mutations were detected but mostly remained low in frequency or were not maintained (Supplemental Table 2 and Supplemental Figure 6). In 19 out of 72 guinea pigs, however, *de novo* mutations that increased in frequency during infection were observed (Supplemental Table 2, Supplemental Figures 6-7). In this latter category, positive selection of the *de novo* variants appeared to drive barcode dynamics, as evidenced by the coordinated sweep of the *de novo* mutation and a single barcode (Supplemental Table 2 and Supplemental Figure 7). Of note, this pattern only occurred in Tx/12 HA F159S BC2; high frequency barcodes in Tx/12 HA WT BC1 were not linked to any *de novo* mutations. Among the *de novo* mutations that showed evidence of positive selection, nearly all were proximal to the HA receptor binding site. One that occurred outside of this region was in the neuraminidase (NA) gene. Another mutation was observed early in infection in the stalk domain of HA. Further quantitative analysis to estimate the number of sites on HA that can compensate for F159S was performed. In the animals with *de novo* sweeps, 37 intra-host single nucleotide variants (iSNVs) above frequency of 10% were observed, including 25 unique sites. Using a Poisson-based coverage equation, the *de novo* mutations identified in the whole genome analysis represent approximately 77% of all compensatory mutations that are estimated to be possible (48 sites in total). Together, these dynamics suggest that compensatory mutations that offset the fitness cost of HA F159S were positively selected within hosts.

### Design of a barcoded antigenic variant virus with eased fitness tradeoffs

Our data suggest a prominent role for fitness tradeoffs in defining the potential for antigenic selection. To test the hypothesis that antigenic selection would proceed readily in their absence, we sought to identify an antigenic variant in which fitness tradeoffs were eased. Analysis of publicly available sequences [35, 36] revealed two mutations that co-occurred with HA F159S in the A/H3N2 lineage: HA A138S and HA N225D (Figure 4A,B). Both residues are proximal to the receptor binding site (Figure 4B).

**Figure 4:**
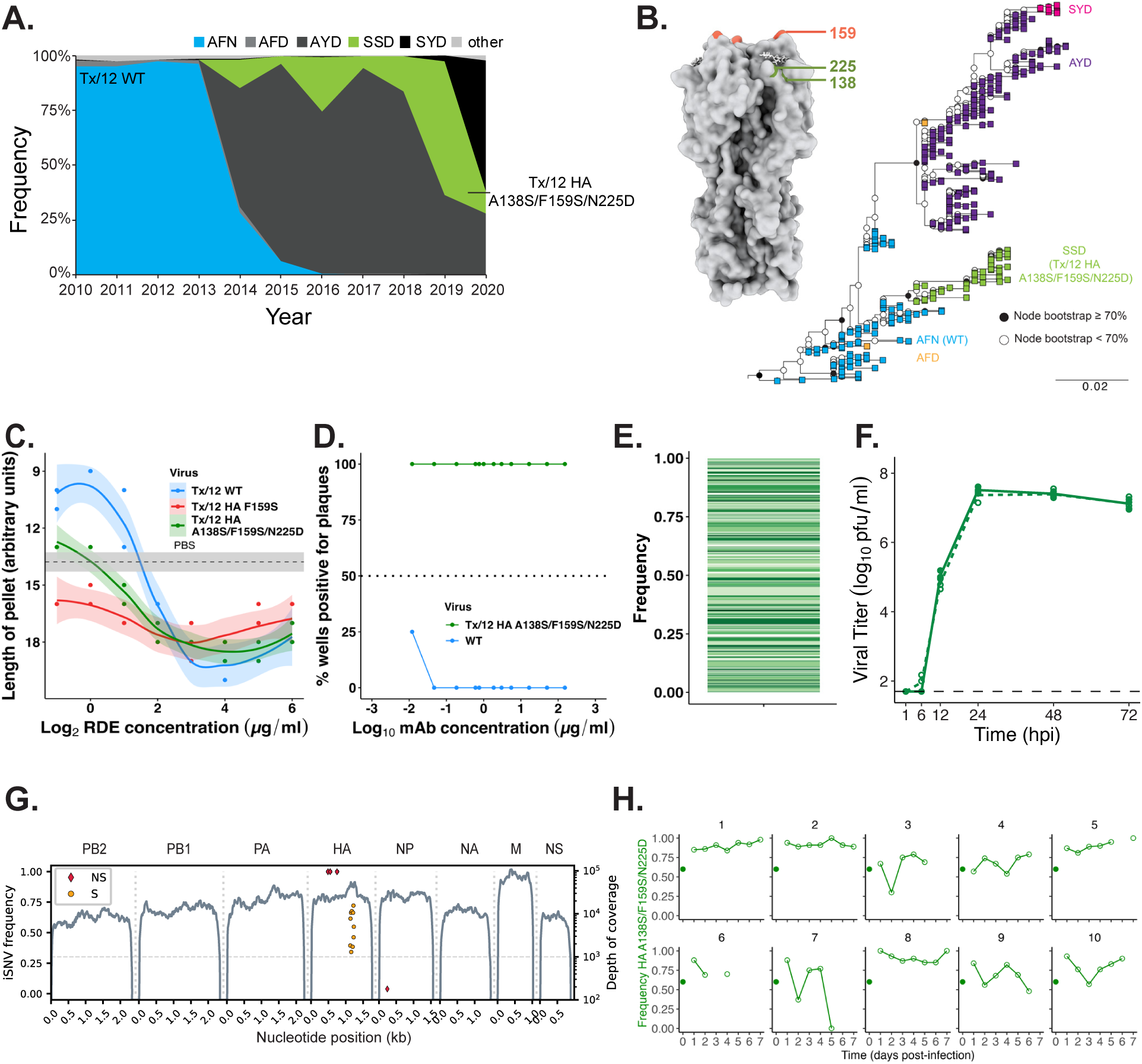
Tx/12 HA A138S/F159S/N225D maintains antigenic escape from mAb 2A05 without detectable fitness costs. **(A)** Area plot showing frequency changes over time of amino acid motifs at HA positions 138, 159, and 225 in the seasonal H3N2 lineage **(B)** Amino acid phylogenetic tree labeled by motifs at HA position 138, 159, 225. HA trimer with positions 138, 159, and 225 labeled (A/X31, PDB: 1HGG). Rooted to A/Iowa/09/2011 (H3N2) HA amino acid sequence. **(C)** Cell binding avidity of Tx/12 WT, HA F159S, and HA A138S/F159S/N225D viruses on receptor destroying enzyme (RDE)-treated red blood cells. Data are expressed as length of red blood cell pellet (performed in duplicate). Solid lines are polynomial regression estimates fit to the data with shaded areas showing 95% confidence intervals. Dashed line represents mean pellet length in PBS negative control with shaded 95% confidence interval. **(D)** Plaque reduction neutralization of Tx/12 HA A138S/F159S/N225D using mAb 2A05. Data are expressed as the percent of wells within each quadruplicate that were positive for ≥1 pfu. Dashed line represents 50% (2/4 replicates) having ≥1 pfu. Tx/12 wild type data from Figure 1 are reproduced for comparison. **(E)** Stacked barplot of barcode frequencies of virus passage 1 stock. Each colored section represents a different barcode, and its height is the relative frequency within the sample. **(F)** Multicycle growth curves of Tx/12 HA A138S/F159S/N225D (green, dashed line) and Tx/12 HA A138S/F159S/N225D BC2 (green, solid line). Infections were performed in triplicate, shown as individual points with the line plotted through the mean. Dashed line represents limit of detection of 50 pfu/ml. Repeated measures ANOVA (p=0.312) was performed. (**G)** Whole genome sequencing of barcoded virus stock. Categorized iSNVs (S=synonymous, NS=nonsynonymous) are plotted at their frequency on the left axis and sequencing coverage (gray line) is plotted on the right axis. Segments are identified above each trace. **(H)** Frequency of the Tx/12 HA A138S/F159S/N225D allele over days of infection when animals were inoculated with ∼50% mutant and 50% wild type viruses. Solid dot denotes starting frequency of Tx/12 HA A138S/F159S/N225D in the inoculum. Guinea pig number is identified above each trace.

To evaluate whether the addition of HA A138S and N225D to the Tx/12 HA F159S background reduced fitness costs of HA F159S, the cell binding avidities of Tx/12 WT, HA F159S, and HA A138S/F159S/N225D viruses were compared. We found that the binding of the triple mutant virus was improved relative to HA F159S alone but was not equivalent to Tx/12 WT virus (Figure 4C). Plaque reduction neutralization assays with mAb 2A05 showed that the Tx/12 HA A138S/F159S/N225D virus maintained an antibody escape phenotype similar to that of the HA F159S virus (Figure 4D).

The triple mutant virus was also outfitted with a genetic barcode and named Tx/12 HA A138S/F159S/N225D BC2. The library had 805 barcodes and a Shannon diversity of 5.6 (Figure 4E). Barcodes did not affect viral growth (Figure 4F) and whole genome sequencing did not reveal *de novo* mutations expected to alter barcode dynamics (Figure 4G).

We assessed the fitness of Tx/12 HA A138S/F159S/N225D BC2 virus relative to Tx/12 HA WT BC1 virus *in vivo* by co-infecting animals with a ∼1:1 mixture of the two viruses (see Supplemental Table 1 for inoculum composition). In all animals, the Tx/12 HA A138S/F159S/N225D BC2 virus predominated over time (Figure 4H). Thus, the addition of A138S and N225D mutations eased the negative fitness effects of F159S, as intended.

### Antigenic selection was more efficient when fitness tradeoffs were lessened

To determine if antigenic selection proceeded efficiently when the fitness cost of the variant was reduced, animals were co-infected with a ∼10:1 mixture Tx/12 HA WT BC1 and Tx/12 HA A138S/F159S/N225D BC2 viruses (see Supplemental Table 1 for inoculum composition). In an experiment in which selection was imposed at the time of infection, animals were given either no mAb or an intermediate dose of mAb 2A05, which was previously seen to be insufficient to drive the selection of the Tx/12 HA F159S BC2 single mutant virus. In addition, in an experiment which assessed the impact of delaying immune pressure, animals were given a high dose of mAb 2A05 starting either at the time of inoculation or 6 h post-inoculation. As before, daily nasal lavage samples were analyzed to define antigenic variant abundance and barcode composition.

In naïve animals, Tx/12 HA A138S/F159S/N225D BC2 remained a minor variant or was lost during infection. However, positive *β* estimates for Tx/12 HA AA138S/F159S/N225D BC2 under this condition indicate an overall increase in frequency (Supplemental Figure 2C). When animals were given an intermediate dose of mAb 2A05, Tx/12 HA A138S/F159S/N225D BC2 was fixed or nearly fixed in all animals (Figure 5A), and *β* was higher than in the absence of immune pressure (Supplemental Figure 2C). This outcome for the triple mutant stands in sharp contrast to the loss or decline of the single mutant under the same conditions (Figure 2A, Supplemental Figure 2A). When a high dose of mAb 2A05 was applied with no delay between inoculation and mAb instillation, Tx/12 HA A138S/F159S/N225D BC2 quickly fixed in all animals (Figure 5A), similar to the single mutant under these conditions (Figure 3A). When this strong immune selection was delayed to 6 h post-inoculation, Tx/12 HA A138S/F159S/N225D BC2 nonetheless increased in frequency and approached fixation in all animals (Figure 5A), yielding *β* estimates > 0 (Supplemental Figure 2C). This robust positive selection of the triple mutant virus under conditions of delayed immune pressure contrasted with the single mutant, where outcomes were varied and *β* estimates were reduced (Figure 3A, Supplemental Figure 2B). Together, these data show that easing fitness tradeoffs increased the efficiency of antigenic selection.

**Figure 5:**
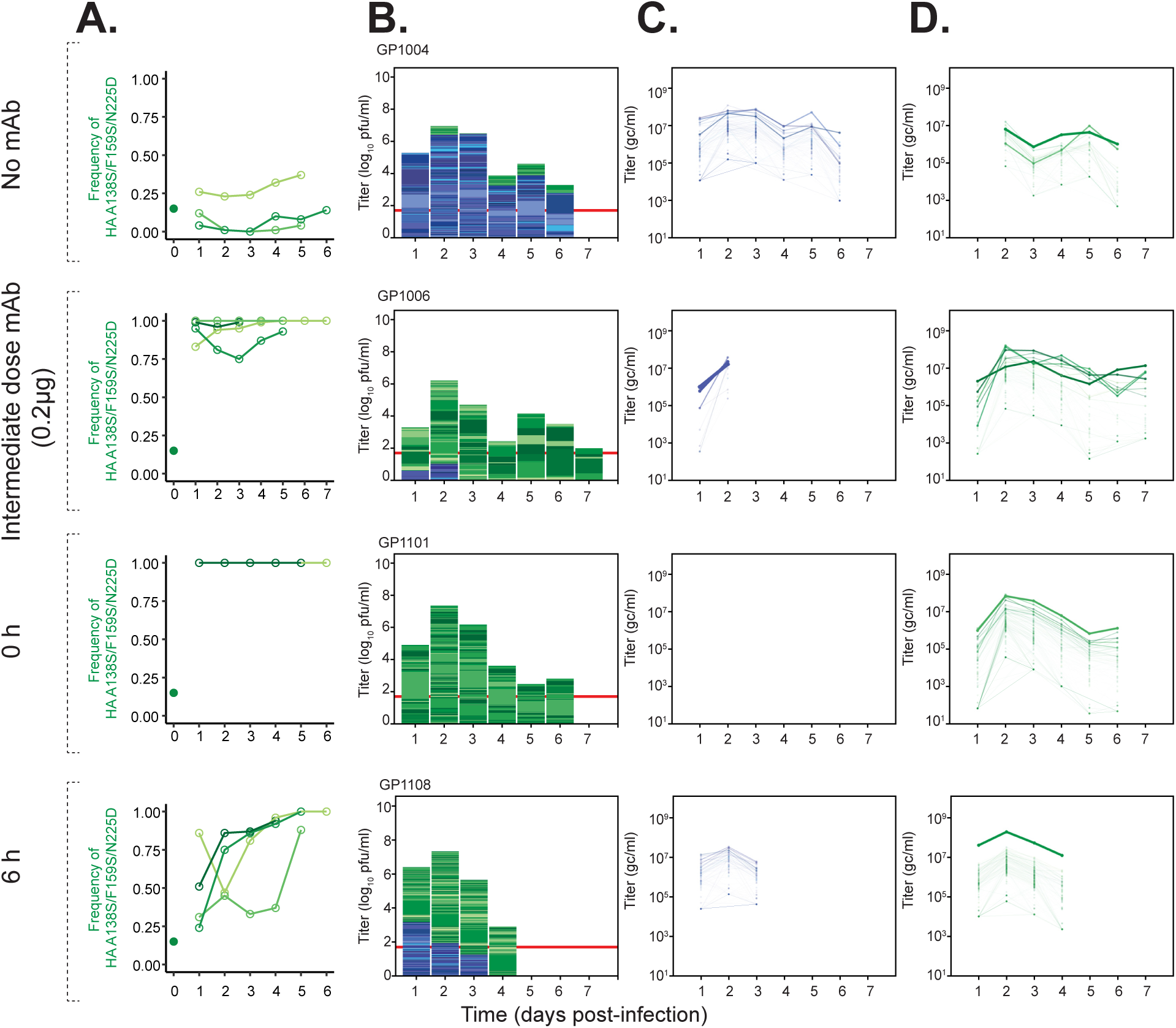
Antigenic selection is efficient when fitness costs are eased through the addition of second-site mutations. Tx/12 HA WT BC1 virus is represented by blue/purple colors and Tx/12 HA A138S/F159S/N225D BC2 is represented by green colors. Animals were inoculated with 90% WT and 10% HA A138S/F159S/N225D BC viruses. At the left of each row, the mAb dose applied at the same time as viral inoculation or the time interval in hours between viral inoculation and the first administration of a high dose of mAb (0.4 ug) is indicated. **(A)** Frequency of the Tx/12 HA A138S/F159S/N225D allele over days of infection. Solid dot denotes starting frequency of Tx/12 HA A138S/F159S/N225D in the inoculum. Each shade of green represents a different animal. Animals in which *de novo* mutations were associated with barcode dynamics are not displayed. Panels **(B)-(D)** contain a representative animal per group (the same animal in all panels). Data from remaining animals are shown in Supplemental Figure 9. **(B)** Stacked bar plots where the total height of the column is scaled to viral titer (pfu/ml). Each colored section within the column represents a unique barcode, and its height is the relative frequency within the sample. The total height of green bars and blue/purple bars is scaled to the gc/ml titer of each virus. Red horizontal line represents limit of detection of plaque assay (50 pfu/ml). **(C)** and **(D)** show Tx/12 HA WT and A138S/F159S/N225D individual barcodes over time, respectively. Each line is a unique barcode with its frequency scaled to the gc/ml titer. Line thickness is scaled to the average frequency of the barcode. Gaps denote results below the limit of detection.

In all conditions, barcode dynamics in these co-infections mirrored those seen with co-infections of the wild type and single mutant viruses. In some animals, many barcodes within the dominant subpopulation were maintained over time, indicating minimal stochastic effects. In other animals, the selection of a subpopulation was driven by only a few barcode lineages, indicating concurrent stochastic effects (Figure 5B-D, Supplemental Figure 8). However, these stochastic effects did not obstruct selection.

### Stochasticity was moderate and driven by variability in high frequency barcodes

Our data indicate that stochastic effects are often muted and, even where they are more prominent, the overall population is nonetheless shaped by selection. To quantify stochasticity, we adopted a method that uses longitudinal frequency dynamics to estimate effective population size (N_e_) [37] and applied this approach to the barcode frequencies observed over time within each viral subpopulation. Co-infected animals were analyzed in this way; importantly, even in the presence of selection distinguishing the two infecting subpopulations, we could still use the barcode frequency dynamics within each subpopulation to infer the N_e_ because the barcodes within a subpopulation are selectively neutral with respect to each other. Animals that were mono-infected with either Tx/12 HA WT BC1 or Tx/12 HA F159S BC2 viruses were additionally used to determine N_e_ in the absence of competition between variant alleles (Supplemental Figure 9). Animals in which *de novo* mutations appeared to drive barcode frequency dynamics were removed from the analysis (Supplemental Figure 7).

In all conditions, we observed that N_e_ was proportional to viral titers and vary by day post-infection, as expected (Figure 6). The average estimate of N_e_ across all conditions was 760, 95% CI [462,1057] at peak viral titer and 594, 95% CI [209, 979] following peak titers, indicative of only moderate stochastic effects as predicted under the Wright-Fisher model (Figure 6) and consistent with the dominance of selective dynamics. Whether in the context of co-infection or mono-infection, and whether in presence or absence of effective immune pressure, N_e_ estimates were comparable among the three different antigenic variant viruses.

**Figure 6:**
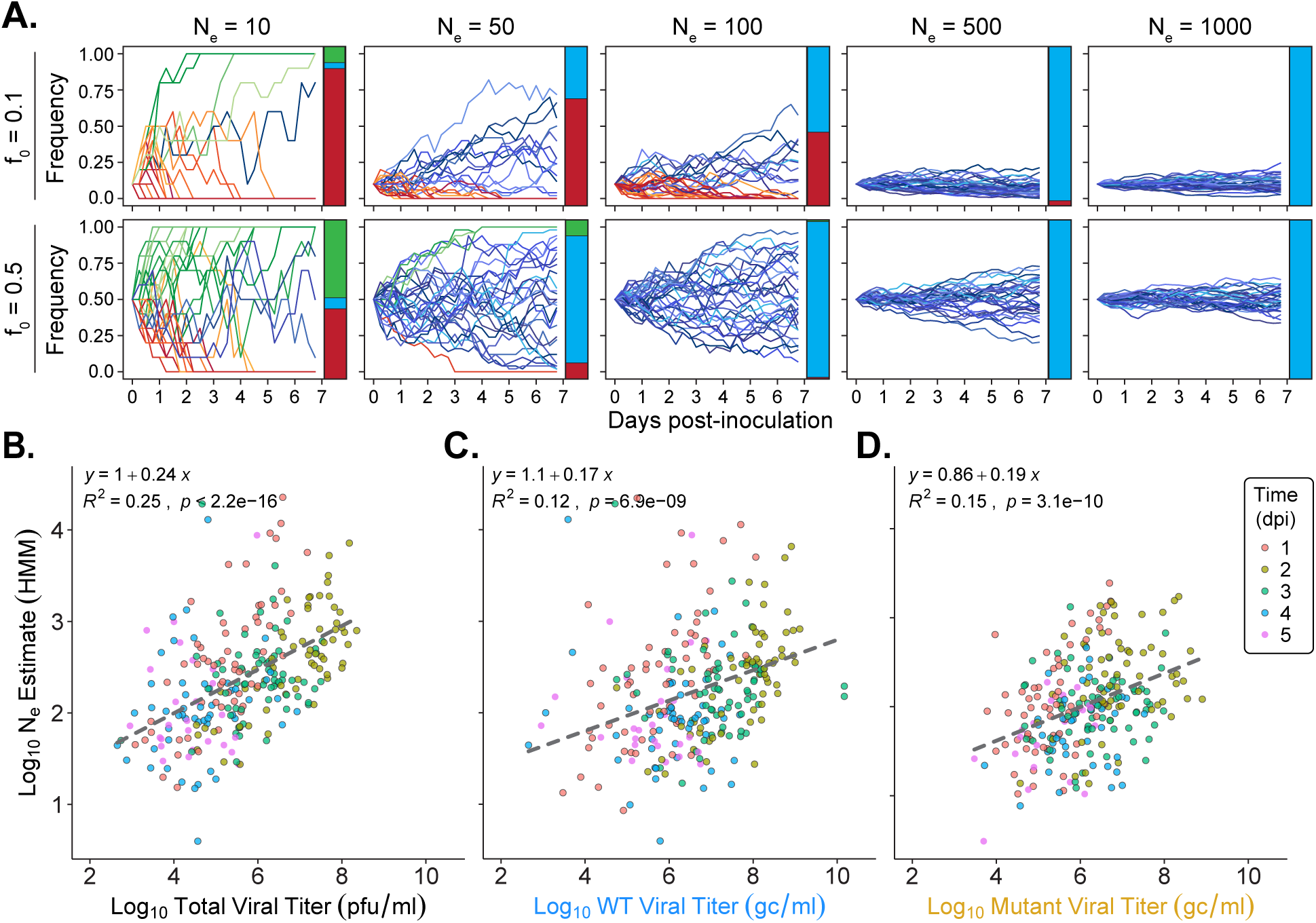
Log N_e_ estimates indicate moderate stochasticity, are proportional to the log of viral titers and vary with day post infection. **(A)** Simulated variant frequency over time under Wright-Fisher assumptions when variant frequency starts at 10% and 50% (f_0_). Different conditions of N_e_ are shown, as indicated above each facet. Each line is the result of one simulation, and a total of 30 simulations were performed. Red colors identify trajectories that go extinct, blue are trajectories that are maintained, and green are trajectories that are fixed. The stacked barplots at the right of each trace show the probability of each trajectory. **(B)** Log N_e_ estimate (using Hidden Markov Model) plotted against log total viral titer (pfu/ml). **(C)** and **(D)** show N_e_ estimates from HA BC1 or HA BC2 regions plotted against the log of wild type or mutant viral titer (gc/ml), respectively. Both HA F159S and HA A138S/F159S/N225D viruses carry HA BC2 and data from these two viruses are therefore combined in panel (D). For panels B-D, data aggregated across all animals and conditions and colored by day post infection (dpi). Dashed line and equation in upper left of each plot denote linear regression (ordinary least squares).

Under the Wright-Fisher model from which N_e_ derives, if a barcode is at frequency *f_t_* at time t and *f_t_*_+1_ at t+1, the change in the square root of the barcode frequency 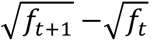 is approximately normally distributed with a standard deviation that is independent of *ft*. Importantly, such a distribution was not observed in our data (Figure 7). Compared to the theoretical expectation, there was an excess of large changes in barcode frequency (Figure 7A) and larger fluctuations in the frequencies of more common barcodes (Figure 7A). This deviation from normality was seen in both co-infected and mono-infected animals and thus was observed both in the presence and absence of selective dynamics (Figure 7B). Furthermore, earlier time points of infection had greater deviation from the expected normal distribution than later time points (Figure 7C). Because high-frequency barcodes showed proportionately more stochasticity than low-frequency ones, estimates of Ne were sensitive to the minimum barcode frequency thresholds used in the analysis, with higher thresholds yielding lower estimates (Figure 7D). These patterns indicate that the Wright-Fisher model is not a good description of the evolutionary dynamics. This may be because the spatial heterogeneity of IAV populations within hosts [32, 38–40] creates environmental stochasticity not captured by the Wright-Fisher model [41].

**Figure 7:**
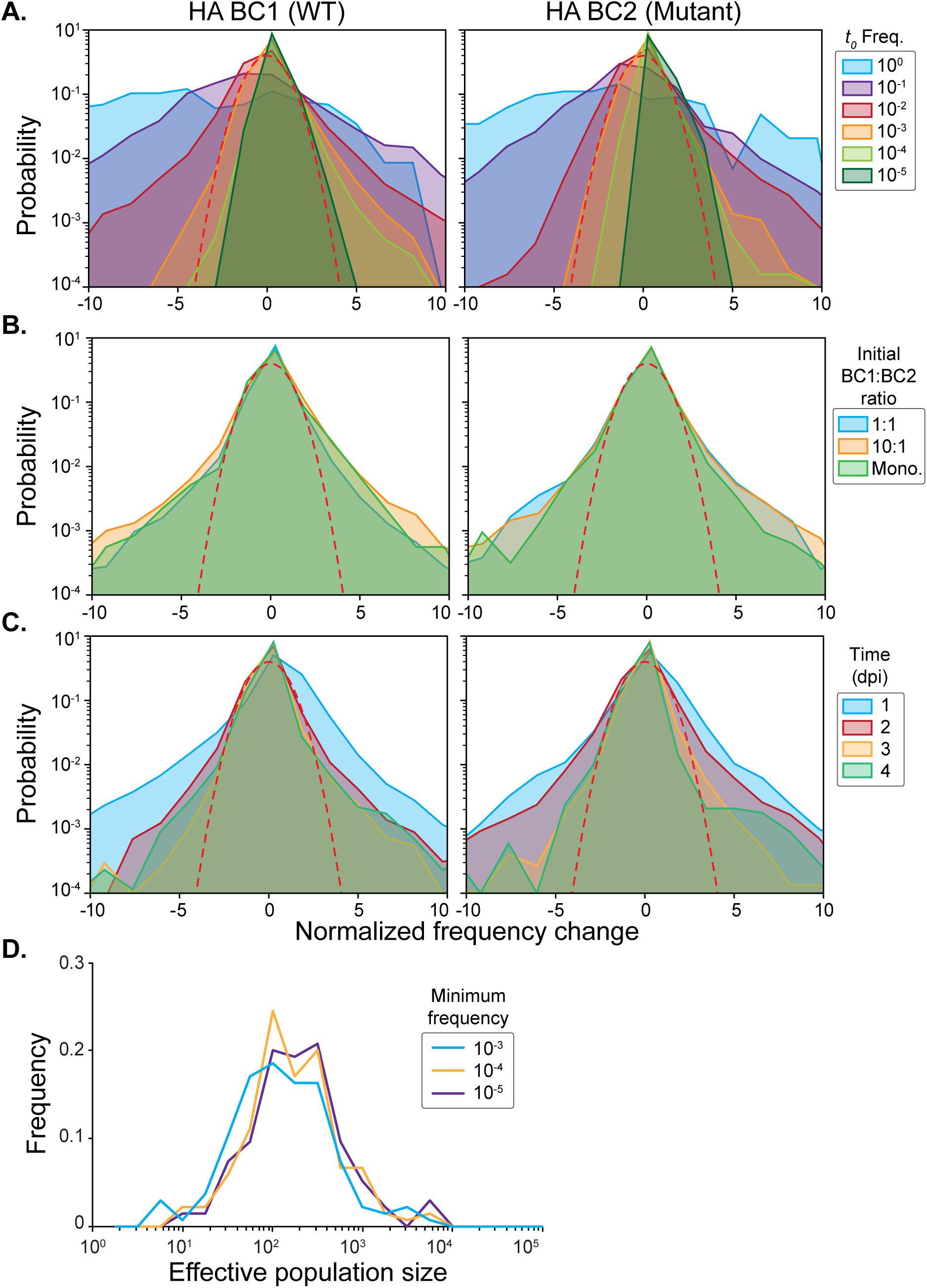
Stochastic effects do not adhere to the Wright-Fisher model of drift. **(A)** Distribution of normalized barcode frequency change colored by starting frequency of the barcode on Day 1. Red dashed outline represents expected distribution predicted under the Wright-Fisher model of drift. Each facet is labeled by barcode region and antigenic variant allele. **(B)** Distribution of normalized barcode frequency change colored by experimental condition. Mono. denotes animals that were mono-infected with either wild type or antigenic variant virus. Red dashed outline represents expected distribution predicted under the Wright-Fisher model of drift. **(C)** Distribution of normalized barcode frequency change colored by timepoint in infection. Red dashed outline represents expected distribution predicted under the Wright-Fisher model of drift. **(D)** Distribution of N_e_ estimates at different minimum frequency thresholds at peak viral titer. Estimates using 10^-5^, 10^-4,^ and 10^-3^ thresholds are shown in different colors.

## DISCUSSION

To understand the potential for seasonal antigenic change to be driven by processes occurring at the within-host scale, we evaluated the dynamics of IAV antigenic selection during acute infection. We found that antigenic selection was limited by fitness tradeoffs and the timing of immune pressure within hosts. Stochastic effects, by contrast, did not present a major barrier to selection. These were moderate, did not adhere to classical predictions of genetic drift, and did not impede selection from ultimately shaping viral populations within hosts.

The observation herein that selection acts efficiently during acute infection contrasts with prior work in naturally infected adults [7–9]. Key factors related to the timing of sample collection may explain why positive selection is not readily observed in these cohorts. The average incubation period for IAV is 3.4 days [42]. Thus, sampling triggered by symptom onset would often miss evolutionary dynamics during the amplification of infection. Thus, selection may have acted early, before sampling was initiated.

Indeed, in our system, selection was most efficient when acting on the incoming viral population during the establishment of infection. This outcome is consistent with the model prediction that immune pressure present at the time of infection enables the fixation of *de novo* antigenic variants [27]. Although not involving antigenic variants, similar dynamics have been previously reported in experimentally infected humans and pigs, with substantial changes in allele frequencies observed early in infection [11, 30, 31]. Based on this observation, it is important to consider that, in natural contexts, the rapidity of antibody waning at the site of infection likely limits the opportunity for the amplification of antigenic variants within hosts [43].

By delaying the introduction of immune pressure, we were able to explicitly model the dynamics of antigenic selection at different stages of antibody recall [23, 24, 44]. We found that delays resulted in loss of the Tx/12 HA F159S variant in a subset of animals, thus eliminating the potential for positive selection of this variant later in infection. Again, this observation is in line with model predictions [27]: recall kinetics in which specific antibody is absent during the virus’ exponential growth phase lead to the loss of *de novo* antigenic variants. In another subset of animals, however, the Tx/12 HA F159S variant was positively selected despite delayed selection. Prior work in ferrets and humans has shown positive selection later in infection [26, 45] and we see this effect in our dataset with the late positive selection of putative compensatory mutations. Thus, where variants are present late, positive selection can act on them. Our data underline the role of fitness tradeoffs in this dynamic: as seen for the Tx/12 HA A138S/F159S/N225D variant, an antigenic variant with little or no fitness cost is more likely to be maintained within-host, extending the opportunity for it to be subjected to immune pressure.

More broadly, two features of our results evidence the importance of functional constraints on HA in defining the likelihood of IAV within-host antigenic evolution. The first is the differing strengths of immune selection needed to favor the Tx/12 HA F159S variant and the Tx/12 HA A138S/F159S/N225D variant. The second is the observed positive selection of *de novo* mutations that may ease the cost of the HA F159S mutation. Prior work has shown that HA antigenic mutations often modulate HA receptor binding, a pleotropic effect that constrains antigenic evolution; epistatic mutations can, however, alleviate this constraint [4, 15, 17–20, 46–48]. In fact, several of the mutations identified in our analyses have been implicated in epistatic networks with residue 159 [15, 18, 49]. Our data build on this work to demonstrate the impact of fitness tradeoffs on antigenic evolution within individual hosts. Our data furthermore show that compensatory mutations arising *de novo* can sweep within the timeframe of an acute infection. This observation is in line with previous reports that describe the sweep *de novo* mutations in experimentally inoculated animals [50–54]. Since our observations were made in the context of an HA F159S mutation, it is notable that this mutation has relatively large negative fitness effects among the set of amino acid changes that have mediated major antigenic transitions in the seasonal H3N2 lineage [55]. Our data explain the rarity with which costly mutations like HA F159S are detected in sub-lineages that drive major antigenic change: the strength of immune selection needed to favor such a variant may be unusual at the within-host level. Importantly, however, our data also indicate that multiple mutational pathways are available to offset the costs of HA F159S, highlighting the need to consider the genetic context of mutations in key antigenic sites when assessing the likelihood of their emergence as major variants.

To vary the timing and strength of immune pressure, we used mucosal application of a monoclonal antibody. It is important to consider how this approach relates to humoral responses in influenza-experienced individuals. The use of a single antibody clone modeled an environment in which a variant carrying a mutation in a single epitope would have maximum fitness gains. Characterization of human serum samples has revealed focused antibody responses to influenza, which ultimately enable antigenic selection [56]. Further, since B cell clonotypes wane at varied rates, a polyclonal response will decline in complexity over time and ultimately become monoclonal [57]. This immune context creates an environment amenable to the amplification of antigenic variants within hosts [57]. Thus, although not broadly representative of human humoral landscapes, the use of a monoclonal antibody in our system recapitulated the natural immunological context in which antigenic selection is most likely to proceed.

We initially hypothesized that stochastic processes acting on within-host viral populations would be a major impediment to selection. On the contrary, our data show that stochasticity across all tested conditions was similar to that of a population experiencing only moderate stochasticity under Wright-Fisher predictions [12, 58]. The average estimates of N_e_ were 760 ± 298 (95% CI) at peak viral titers and 594 ± 385 (95% CI) at later time points. These estimates reflect the continuity of barcode composition seen over time in many of the animals and are consistent with our prior observations in guinea pigs [21]. Results reported to date from naturally infected humans differ, however, in that stochastic effects appear to be prominent [7–10, 13] and N_e_ estimates are in the range of 50-300 [13, 14]. N_e_ was also found to be low in humans and ferrets inoculated experimentally [31], suggesting that the difference is not a consequence of the mode of viral introduction. The difference may stem from the low diversity of typical within-host viral populations [7–10, 13, 31] and the technical challenges this presents for evaluating stochastic effects. Specifically, when putatively neutral variants are used [13, 14], strict frequency thresholds must be applied; our results suggest that this may cause N_e_ to be underestimated. And when trajectories of selected alleles are used [31], it is difficult to distinguish between fluctuations in selection and the true demographic stochasticity described by N_e_. By contrast, the viral barcoding approach used herein enables confident detection of frequency changes across hundreds of neutral variants simultaneously, supporting robust quantification of stochastic effects.

While estimating N_e_ offers a practical means of quantifying the impact of stochastic dynamics on allele frequencies, this approach assumes that the population under study behaves according to the Wright-Fisher model of genetic drift [12, 58, 59]. The patterns of allele frequency changes in our data indicate that these assumptions do not hold. Namely, compared to the Wright-Fisher model [12, 58, 59], we observed an excess of anomalously large frequency changes and high frequency barcodes experienced greater than expected stochasticity, while low frequency barcodes experienced lower than expected stochasticity. This effect was more pronounced at early stages of infection and occurred in both co-infected and mono-infected animals where Tx/12 antigenic alleles were not in competition. Together, this implies that viral processes beyond selection and occurring early in infection drove deviations from Wright-Fisher dynamics. A key assumption of the Wright-Fisher model is that populations are well mixed, such that variation in progeny output is even across the population. On the contrary, within-host influenza virus populations show spatial structure [32, 38–40]. Thus, the relatively high stochasticity observed for higher frequency barcodes may stem from spatial variation in viral growth rates, which in turn is likely to vary over time as the viral population expands to new regions of the respiratory tract. Several processes common in early viral infection may contribute to these dynamics. Viral burst size from single cells is highly variable, especially under conditions of low multiplicity [60–63]. Innate immune signaling is also heterogenous at the single cell level [64, 65], and the distribution of permissive cell types is variable throughout the upper respiratory tract [66]. Combined with superinfection exclusion [67, 68], this anatomical landscape likely creates extensive heterogeneity in the availability of target cells, creating a patchwork of foci with varying growth rates. While formal analysis is needed to test whether these biological processes can account for the barcode frequency dynamics observed herein, our data motivate detailed examination of the sources of stochasticity shaping within-host viral populations.

Our data provide insight into the rarity with which antigenic selection is observed in individual hosts. However, opportunity for antigenic selection extends beyond individuals, to transmission between hosts. Prior work has shown that transmission is associated with a sharp drop in viral diversity [7, 21, 22, 25–29]. Depending on the virus-host context, this contraction of diversity has been attributed to stochasticity [7, 21, 22, 25] or selection [10, 26–29]. The positive selection of antigenic variants may therefore occur during transmission, supplementing instances of within-host antigenic selection [27]. However, further work is needed to examine the role that transmission plays in antigenic evolution.

There are limitations to our research that are important to consider. The intranasal instillation of a relatively large viral population differs from natural infection in both the initial population size and its spatial distribution. Our model therefore does not reflect the earliest population dynamics that follow transmission. In addition, the intranasal instillation of monoclonal antibody may not model the spatial distribution of antibodies in an authentic response and variation in local antibody concentrations may be an important factor shaping within-host antigenic selection. Furthermore, owing to the use of a human IgG in guinea pigs, Fc mediated functions such as antibody-dependent cellular cytotoxicity were absent from our system. Finally, an anti-antibody response to this foreign protein may have weakened immune pressure at later times. We expect this last effect to be minimal in the timeframe of our experiments, however, since serum antibodies are typically detected around 7 days post-antigen exposure [69, 70].

In summary, our analyses showed that the effects of positive selection on within-host IAV populations are readily apparent; however, opportunity for the positive selection of antigenic variants is limited by fitness tradeoffs and the timing of immune selection. Importantly, stochastic effects were moderate and did not prevent selection from driving the evolution of within-host viral populations. Where stochastic effects were observed, the resulting dynamics furthermore did not adhere to the standard model of genetic drift. Together, these data offer valuable mechanistic insight into the forces shaping the fate of antigenic variants within hosts, enabling deeper understanding of global patterns of influenza virus antigenic evolution.

## ACKNOWLEDGEMENTS

We thank Scott Hensley and Patrick Wilson for sharing the plasmids to express mAb 2A05. This work was funded in part by the National Institutes of Health through R01AI165644 and in part by Emory University funds awarded to A.C.L., as well as by the National Science Foundation through PHY-2146260 by D.B.W. V.R. was supported by NIAID through F31AI179207 and by the Atlanta Chapter ARCS Foundation through a Herz Global Impact Award. This study was supported in part by the Emory Integrated Genomics Core (EIGC; RRID:SCR_023529), which is subsidized by the Emory University School of Medicine and is one of the Emory Integrated Core Facilities. Additional support was provided by the Georgia Clinical & Translational Science Alliance of the National Institutes of Health under Award Number UL1TR002378. The content is solely the responsibility of the authors and does not necessarily reflect the official views of the National Institutes of Health. The Emory NPRC Genomics Core (RRID:SCR_026418) is supported in part by NIH P51 OD011132. Sequencing data was acquired on an Illumina NovaSeq 6000 funded by NIH S10 OD026799.

## AUTHOR CONTRIBUTIONS

Conceptualization, V.R. and A.C.L.; data collection, V.R., D.V., C.L., L.O., N.V.M., M.G.; data analysis, V.R., D.V., G.A.B., and D.B.W.; provision of key reagents, J.W.; preparation of initial manuscript draft, V.R., D.V., and A.C.L.; review and editing of manuscript, all authors; funding acquisition, A.C.L.

## SUPPORTING INFORMATION

Document S1. Figures S1-S5, Figures S7-S9, Tables S1 and S2 Document S2. Figure S6

Table S3. Barcode mutations, related to Figure 1A Table S4. Oligonucleotides list

## DATA AVAILABILITY

Viral sequencing data have been deposited at NCBI’s Short Read Archive at SRA: PRJNA1494584 and will be publicly available upon publication.

Viral titer and variant frequency data will be publicly available upon publication at FigShare: https://doi.org/10.6084/m9.figshare.33043946

All original code and versions used has been deposited at FigShare: https://doi.org/10.6084/m9.figshare.33043946

The latest version of pipeline for whole genome analysis is deposited at GitHub: https://github.com/Lowen-Lab/FluSAP

The latest version of pipeline for barcode sequencing analysis is deposited at GitHub: https://github.com/Lowen-Lab/BarcodeID

## MATERIALS AND METHODS

### Ethical considerations

All animal experiments were conducted in accordance with the Guide for the Care and Use of Laboratory Animals of the National Institutes of Health. The studies were conducted under animal biosafety level 2 containment and approved by the IACUC of Emory University (PROTO201700595) for the guinea pig (*Cavia porcellus*). The animals were humanely euthanized following guidelines approved by the American Veterinary Medical Association.

### Guinea pigs

Outbred Hartley strain guinea pigs were obtained from Charles River Laboratories (051). Male and female animals weighing 300-350 g and aged 4-5 weeks old were used for all studies. Influenza viruses transmit between co-housed guinea pigs [71]; thus, to ensure that individual animals can be treated as independent experimental replicates, they were singly housed starting at the time of inoculation and for the duration of the experiment. Male and female guinea pigs were randomly assigned to different treatment groups. A total of 125 animals were used. All procedures and animal care conformed to the guidelines set forth by Guide for the Care and Use of Laboratory Animals of the National Institutes of Health, American Veterinary Medical Association, and with the approval of the IACUC of Emory University.

### Cell lines and maintenance

Madin–Darby canine kidney (MDCK) cells were a gift from Dr. Daniel Perez, University of Georgia, Athens, GA. A seed stock of MDCK cells at passage 23 was amplified and maintained in Minimal Essential Medium (Gibco) supplemented with 10% fetal bovine serum (FBS; Atlanta Biologicals) and Normocin (Invivogen). 293T cells (ATCC, CRL-3216) were maintained in Dulbecco’s Minimal Essential Medium (Gibco) supplemented with 10% FBS and Normocin. All cells were cultured at 37 °C and 5% CO_2_ in a humidified incubator. The cell lines were not authenticated. All cell lines were tested monthly for Mycoplasma contamination while in use. The medium for the culture of influenza A virus in MDCK cells (virus medium) was prepared by supplementing Minimal Essential Medium with 4.3% bovine serum albumin (BSA; Sigma-Aldrich) and Normocin.

### Monoclonal antibody generation and storage

The mAb 2A05 expression and purification were done as previously described [33, 72, 73]. Briefly, the mAb 2A05 heavy and light variable chains (V_H_ and V_L_) were cloned into human IgG expression vectors. mAb 2A05 was produced by transfecting 293T cells with both heavy and light chain plasmids and purified using protein A/G magnetic beads. The mAb 2A05 was then aliquoted and stored at 4°C for short-term storage or flash-frozen in a dry ice and alcohol slurry and stored at -80°C for long-term storage. Flash-freezing did not alter mAb neutralization activity as determined by plaque reduction neutralization assay (see below).

### Generation of Tx/12 HA BC plasmids

The regions of Tx/12 HA in which the two barcode regions were to be inserted were identified by aligning 545 sequences of H3N2-subtype influenza A viruses from 2003 through 2020 with no geographical restrictions. Numbering of nucleotide positions of this alignment used the beginning of the 5’ UTR of the positive sense HA segment a position 1. Two regions which carried substantial synonymous variation were identified at nucleotide positions 306-404 and 1131-1235. Ten synonymous mutations were selected within each region to be used for creating barcode 1 and barcode 2, respectively (Table S3). Based on the identified mutations, double-stranded Ultramers (IDT) containing degenerate bases with two naturally occurring nucleotides at each of the 10 chosen barcode sites were designed (Tx12_HABC1_ultramerF/R and Tx12_HABC2_ultramerF/R).

Prior to insertion of the Ultramers encoding the barcodes, an XhoI restriction site and two stop codons were introduced within each barcode region in the wild type reverse genetics plasmid, pDP Tx/12 HA, by site directed mutagenesis (QuikChange, Agilent). The three mutations in each region were added successively (using Tx12_HABC1_XhoIF/R, Tx12_HABC1_stop1F/R, Tx12_HABC1_stop2F/R, Tx12_HABC2_XhoIF/R, Tx12_HABC2_stop1F/R, and Tx12_HABC2_stop2F/R primers). Thermocycler conditions were 95°C for 30 s followed by 16 cycles of 95°C for 30 s, 55°C for 1 min, and 68°C for 4.5 min then 68°C for 5 min and hold at 10°C. PCR products were digested with DpnI (NEB, 37°C for 16 h) to remove unmodified plasmid DNA. Introduction of an XhoI site allowed subsequent restriction digestion to destroy residual parental template, while the inclusion of stop codons rendered any remaining wild type pDP Tx/12 HA nonfunctional. Both types of modifications were used to minimize wild type carry-though in the cloning and virus generation steps. Modifications were confirmed by Sanger sequencing (using Tx12_HABC1_78F and Tx12_HABC2_1019F) after each round of mutagenesis following transformation in DH5ɑ cells (Agilent) and plasmid purification (Qiagen QIAprep Spin Miniprep Kit).

To generate a linearized template for barcode insertion, the following steps were performed: XhoI (NEB) digestion of 1 µg of the modified plasmid stock (37°C, 1 h and hold at 10°C) was followed by phosphatase treatment by rSAP (NEB) (37°C, 45 min and 65°C for 20 min, hold at 10°C) to dephosphorylate cut ends of the plasmid. The plasmid was then amplified by PCR (Agilent) using primers that extended outwards from each barcode region: Tx12_HABC1_400down and Tx12_HABC1_298up (95°C for 2 min, then 30 cycles of 95°C for 30 s, 52°C for 30 s, 72°C for 5 min, followed by 72°C for 10 min and hold at 10°C) and Tx12_HABC2_1234down and Tx12_HABC2_1123up (95°C for 2 min, then 30 cycles of 95°C for 30 s, 50°C for 30 s, 72°C for 5 min, followed by 72°C for 10 min and hold at 10°C). Each reaction was performed in duplicate. Duplicates were combined and the linearized product was isolated using column purification (Qiagen QIAquick PCR Purification Kit). To remove residual wild type plasmid, 2 µg of linearized product was subjected to double digestion with DpnI and XhoI (37°C for 90 min) and phosphatase treatment with rSAP (37°C for 30 min, 80°C for 20 min, cool down by 10°C steps every 30 s until hold at 10°C). Column purification was repeated followed by an assembly reaction using the NEBuilder HiFi DNA Assembly Kit (NEB) to insert the Ultramers into the linearized vector and re-circularize (50°C for 1 h, 4°C hold). The product was then transformed into DH5ɑ cells (NEB) in the following manner: 8 transformations for each barcode region were performed using 1.5 µl of assembly product and 100 µl of DH5ɑ cells according to manufacturer protocol. Following recovery incubation, the 8 transformations were pooled together for each construct and centrifuged for 5 min, 2000 RPM, room temperature (Sorvall ST 16). The bacterial pellet was resuspended in 1 ml of 2XYT broth and plated across 5 LB-ampicillin plates in equal volume and incubated for 16 h at 37°C. Approximately 1x10^4^ colonies were collected and pooled in LB-ampicillin culture media and incubated at 37°C for 5 h shaken at 220 RPM. The plasmid was purified from the bacterial population by maxiprep (Qiagen Plasmid Maxi Kit). The presence of a mixed nucleotide identifies within the barcode region in the plasmid stock was verified by Sanger sequencing (using Tx12_HABC1_78F and Tx12_HABC2_1019F). These plasmid stocks were then used to generate Tx/12 HA WT BC1, Tx/12 HA F159S BC2, and Tx/12 HA A138S/F159S/N225D BC2 viruses by reverse genetics in combination with seven plasmids encoding the remaining Tx/12 WT gene segments in a pDP2002 vector [74].

### Viruses

Tx/12 non-barcoded and barcoded viruses were generated by reverse genetics. 293T cells (2.5x10^5^ cells/ml) and MDCK cells (4.5x10^5^ cell/ml) were co-cultured in OptiMEM medium (Thermo Fisher) and 0.18% Normocin for 24 h in a six-well plate. Eight reverse genetics plasmids (1 µg each) were combined with 16 µl of TransIT-293 reagent (Mirius) diluted in 104 µl of OptiMEM and incubated for 1 h at room temperature followed by the addition of 800 µl of OptiMEM. Media was removed from the co-cultured cells and the transfection complexes were added and incubated for 24 h at 37°C. Following incubation, 1 ml of OptiMEM, 0.18% Normocin, and 2X tosyl phenylalanyl chloromethyl ketone (TPCK) treated trypsin (Sigma-Aldrich) were added to each well and incubated for 48 h at 37°C. The medium was then collected and spun at 2500 RPM for 5 min to pellet cell debris. The supernatant was aliquoted and stored at -80°C. Plaque assays were performed to titer virus present in the supernatant (see below).

Recovered virus was propagated in MDCK cells to generate working stocks. In brief, 4-5 T75 flasks containing ∼10^7^ cells per flask were washed three times with 5 ml of 1X PBS (Corning) each and were infected at an MOI of 0.01 pfu/cell in 1X PBS. This MOI was chosen to avoid accumulation of defective viral genomes but seed a sufficient viral population size to maintain barcode diversity. Following a 1 h incubation at 37°C, virus medium supplemented with 1X TPCK trypsin was added, and flasks were incubated for 42-45 h at 37°C. The media from the flasks for each stock were pooled and centrifuged for 1000 RPM for 5 min (Sorvall ST 16). Clarified supernatant was aliquoted and stored at -80°C. Viral titers were determined by plaque assay in MDCK cells and the presence of a diverse barcode in the virus stock was verified by next-generation sequencing (see below).

### Multicycle growth curves

Growth kinetics of Tx/12 barcoded and non-barcoded viruses were determined in triplicate culture wells. MDCK cells were seeded at 2x10^5^ cells/ml in six-well dishes 24 h prior to infection. Cells were washed 3 times with 1X PBS and inoculated at an MOI of 0.01 pfu/cell in 1X PBS. After 1 h incubation at 37°C, inoculum was removed, cells were washed 3 times with 1X PBS, 2 ml of virus medium supplemented with 1X TPCK trypsin was added to cells, and dishes were returned to 37°C. A 120 µl volume of culture medium was sampled at the indicated time points and aliquoted and stored at -80°C. Viral titers were determined by plaque assay on MDCK cells (see below). Plaque assay titers of Tx/12 barcoded and non-barcoded viruses were evaluated by repeated measures ANOVA using the rstatix package version 0.7.2 and plotted using the tidyverse package version 2.0.0 in RStudio.

### Plaque reduction neutralization assay

Plaque reduction neutralization titers were determined in quadruplicate culture wells. MDCK cells were seeded 24 h prior to infection in 24-well culture plates. Tx/12 non-barcoded viruses were diluted to 100 pfu/50 µl and mAb 2A05 was diluted from 300 µg/ml to 0.024 µg/ml in 3-fold increments in 1X PBS. Equal volumes of virus and mAb 2A05 were mixed for a final concentration of mAb 2A05 ranging from 150 µg/ml to 0.012 µg/ml. A negative control was included by mixing virus and 1X PBS. Mixtures were incubated for 1 h at 37°C. Following incubation, cells were washed once with 1X PBS and inoculated with 60 µl of virus-mAb mix. Cells were incubated for 1 h at 37°C with occasional rocking, after which the inoculum was removed, cells were washed once with 1X PBS, and overlayed with 1 mL of plaque assay medium (2X Minimal Essential Media (Gibco) supplemented with 4 mM L-glutamine (Gibco), 30 mM HEPES buffer (Corning), 2X Penicillin-Streptomycin Solution (Corning), 0.3% sodium bicarbonate (Corning), 0.42% bovine serum albumin (BSA) (Sigma-Aldrich), 0.01% diethylaminoethyl (DEAE)-dextran (MP Biomedicals), 1 μg/mL TPCK-treated trypsin and diluted to a 1X concentration with the addition of cell culture grade water (Corning) and a final concentration of 0.7% Oxoid agar (Oxoid) melted and then cooled to 55°C). Plates were incubated for 48 h at 37°C. After incubation, cells were fixed with 1 ml of 10% neutral buffered formalin (VWR) incubating for 1 h at room temperature followed by staining with crystal violet solution (Sigma-Aldrich) to identify any wells that had ≥1 pfu. Neutralization of Tx/12 non-barcoded viruses by mAb 2A05 was calculated by determining the percentage of wells (out of 4) in which ≥1 pfu was detected across mAb 2A05 concentrations and plotted using the tidyverse package version 2.0.0 in RStudio.

### Cell binding avidity assay

Cell binding avidity was evaluated in duplicate for each virus. Receptor destroying enzyme (RDE; Sigma-Aldrich, #C8772) was resuspended in sterile cell culture grade water (Corning) [75]. RDE was further diluted in sterile calcium saline solution (0.1% w/v calcium chloride dihydrate [Fisher], 0.9% w/v anhydrous sodium chloride [Fisher], 0.12% w/v anhydrous boric acid [VWR], and 0.0052% sodium borate decahydrate [ThermoScientific], pH 7.2) [75] and stored in single use aliquots at -80°C. Turkey erythrocytes (Lampire Biological Laboratories) were washed and prepared in a 4% (vol/vol) mixture in 1X PBS. RDE aliquots were thawed and diluted 0.5-64 µg/ml in calcium saline and combined in equal parts with the turkey erythrocyte suspension. A negative control of turkey erythrocytes in calcium saline alone was included. The mixtures were incubated for 1 h at 37°C with occasional mixing following which treated erythrocytes were washed in 1X PBS. Washed erythrocytes were resuspended in 2% (vol/vol) solution in 1X PBS. Viruses were diluted to 4 agglutinating doses (determined by agglutination on untreated turkey erythrocytes) and 12.5 µl of each treated erythrocyte solution was added to 50 µl of diluted virus in a 96-well v-bottom plate. Negative controls consisted of 50 µl diluted virus and 12.5 µl of 1X PBS. Mixtures were allowed to incubate for 1 h at room temperature following which the plate was tilted 45° for 5 min and an image was captured. Data are expressed as maximum concentration of RDE that still enabled agglutination as measured by the cell pellet length. Erythrocyte pellet length was measured by ImageJ (version 2.3.0/1.53q) [76] and plotted across RDE treatment concentrations using the tidyverse package version 2.0.0 in RStudio.

### Guinea pig antibody instillation and infection

Male and female Hartley strain guinea pigs weighing 300-350 g were obtained from Charles River Laboratories and housed by Emory University Division of Animal Resources. Guinea pigs were sedated with ketamine (40 mg/kg) and xylazine (2 mg/kg) by intramuscular injection prior to intranasal inoculation, intranasal antibody instillation, or nasal lavage. The mAb 2A05 was given intranasally in a 100 µl volume of PBS at required amounts (measured in µg of mAb 2A05). Virus inoculum was given intranasally in a 30 µl volume of PBS containing 5x10^4^ pfu of Tx/12 BC viruses. Nasal lavage was performed with 1 ml of PBS per animal on days 1-7 post-inoculation. Collected fluid was divided into aliquots and stored at -80°C. mAb 2A05 was re-instilled as described above following nasal lavage on days 1-7 post-inoculation. Viral titers of nasal lavage samples were determined by plaque assay following which viral RNA was extracted to perform genotypic assays (see below).

### Plaque assay

MDCK cells were seeded 24-28 h prior to infection in 6-well culture plates. Samples were diluted 10-fold from 10^-1^ to 10^-6^ in 1X PBS. PBS-washed cells were inoculated with 200 µl of diluted virus sample and incubated for 1 h at 37°C with occasional rocking. Inoculum was removed and cells were washed again with 1X PBS, following which cells were overlayed with 2 mL of plaque assay medium as described above. Plates were incubated for 48 h at 37°C. Plaques were counted by visual inspection under oblique lighting.

### Viral RNA extraction

Viral RNA was extracted from nasal lavage or inoculum samples using either the QiaAmp Viral RNA kit (Qiagen) or the NucleoMag RNA extraction kit (Macherey-Nagel). QiaAmp extractions followed manufacturer guidelines except that carrier RNA was omitted and final elution used 30 µl of nuclease-free water. NucleoMag extractions used the EpMotion liquid handler (Eppendorf) and followed manufacturer guidelines except that rDNase was omitted and final elution used in 47 µl of nuclease-free water.

### Droplet digital PCR (ddPCR) assay

cDNA was synthesized from 12 µl of RNA in 20 µl reactions using Maxima reverse transcriptase (Thermo Scientific) with the provided buffer, dNTPs at a final concentration of 0.5 mM, Ribolock RNase inhibitor (Thermo Scientific), and UnivF(G)+6 and UnivF(A)+6 primers[21, 77] at a final concentration of 0.15 µM each. Samples were incubated at 55°C for 30 min, 85°C for 10 min, held at 10°C temporarily and stored at -20°C.

DdPCR was performed using the QX200 Droplet Digital PCR system (Bio-Rad) on samples that were positive by plaque assay. Each reaction included ddPCR Supermix (no dUTP) (Bio-Rad), Tx12_ddPCR_primerF/R (IDT) at a final concentration of 0.1 µM each, Tx12_ddPCR_WT_probe and Tx12_ddPCR_HAF159S_probe (IDT) at a final concentration of 0.5 µM each, and diluted cDNA. Droplets were generated according to the manufacturer’s instructions [78] and transferred to a 96-well PCR plate. Plates were heat sealed, and targets were amplified using the following thermocycling conditions with all ramp rates set to 2°C/s: 95°C for 10 min, 49 cycles of 94°C for 30 s and 62.6°C for 1 min, 98°C for 10 min, and hold at 4°C. Samples were then analyzed on the QX200 Droplet Reader using QuantaSoft software (Bio-Rad, version 1.7.4.0917), and fluorescence thresholds set manually based on negative controls.

### Targeted barcode sequencing

Targeted barcode sequencing was performed on all samples positive in plaque assay. The barcoded regions of Tx/12 were amplified using the Superscript III One-step RT-PCR with Platinum Taq High Fidelity DNA polymerase (Invitrogen) and the following primers [79]: Tx12_HABC1seq_F, Tx12_HABC1seq_R, Tx12_HABC2seq_F, and Tx12_HABC2seq_R (IDT) each at 0.4 µM final concentration. Thermocycling conditions were 55°C for 2 min; 45°C for 60 min; 94°C for 2 min; 30 cycles of 94°C for 30 s, 59.2°C for 30 s, 68°C for 1 min; 68°C for 5 min, and hold at 10°C. DNA fragments were purified using NucleoMag NGS Clean-up and Size Select (Macherey-Nagel) at a 2.5x (vol/vol) bead to DNA preparation ratio. Quantification of purified DNA was done using Qubit dsDNA high sensitivity kit (Thermo Fisher). Library preparation (Nextera XT, Illumina), sequencing (MiSeq, 2x150 bp paired end reads, Illumina), and demultiplexing was performed at the Emory Integrated Genomics Core facility.

### Analysis of barcode sequencing data

Barcode frequencies and diversity metrics were determined using BarcodeID (see FigShare and GitHub links) as previously described [21, 32, 40, 80]. Shannon diversity was calculated according to the formula *H* = −∑*p_i_* ln(*p_i_*) where *pi* is the frequency of the barcode *i*.

### Viral whole genome sequencing

Viral whole genome sequencing was performed on a subset of samples that were positive by plaque assay. Multi-segment PCR amplification of the viral genome was performed using 2.5 µl of purified RNA, Uni/Inf primers [77], and the Superscript III One-step RT-PCR with Platinum Taq High Fidelity DNA polymerase. The thermocycler amplification protocol was 55 °C for 2 min, 45 °C for 60 min, 94 °C for 2 min, then 5 cycles of 94 °C for 30s, 44 °C for 30 s, 68 °C for 3.5 min, followed by 26 cycles of 94 °C for 30s, 57°C for 30s and 68°C for 3.5 min, then 68°C for 10 min and lastly, hold at 4°C. DNA fragments were purified using NucleoMag NGS Clean-up and Size Select at a 0.45x (vol/vol) bead to DNA preparation ratio. Quantification of purified DNA was done using Qubit dsDNA high sensitivity kit. Library preparation, sequencing (NovaSeq 6000, 2x100 bp paired end reads, Illumina), and demultiplexing was performed at the Emory National Primate Research Center Genomics Core facility.

### Analysis of whole genome sequencing data

Whole genome sequencing reads were analyzed using the FluSAP pipeline (see FigShare and GitHub links) as previously described [21, 80–82] with the following modifications: minor variants were required to be present at ≥3% frequency, reads containing the minor allele needed an average mapping quality score of ≥30, the reads overall needed to have a ≤10.0 average mismatch and ≤1.0 indel counts relative to the reference sequence, and putative iSNVs were required to have a frequency at least three standard deviations above the background distribution of minor allele frequencies in the surrounding 100 nucleotides, and only sites with ≥1000x coverage were considered.

### Verification of inoculum composition ratios

To validate the ratios of variant viruses present in inoculums used for co-infection of guinea pigs, we isolated 48 plaques from each mixture and determined their genotypes using sequence-specific probes in ddPCR or qPCR, as follows.

MDCK cells were seeded on 10 cm dishes 24-28 h prior to infection with 4-6 dishes seeded per inoculum. Medium was removed and cells were washed once with 1X PBS. Inoculum was diluted to 100 pfu/ml and 200 pfu/ml, and cells were inoculated with 1 ml of diluted inoculum followed by incubation for 1 h at 37°C with occasional rocking. Following incubation, inoculum was removed, cells were washed once with 1X PBS, and 12 ml of agar overlay was added per dish as described previously. Dishes were incubated for 48 h at 37°C.

Following incubation, 48 well-separated plaques across the dishes were isolated using a 5 mL serological pipette and transferred to a 96-well block containing 160 µl of 1X PBS per well and stored at -80°C. Viral RNA was extracted using the NucleoMag RNA extraction kit as described previously. Plaque-derived viral RNA was then either used to generate cDNA and genotyped via ddPCR (as described previously) or used directly with the iTaq Universal Probes One-Step Kit (Bio-Rad) and the CFX384 Real-Time System (Bio-Rad) in the following manner: each 20 µl reaction included the iTaq Rx Mix and iScript Rev Pol according to manufacturer guidelines, Tx12_ddPCR_primerF/R (IDT) at a final concentration of 450 nM each, Tx12_ddPCR_WT_probe and Tx12_ddPCR_HAF159S_probe (IDT) at a final concentration of 250 nM each, and 3 µl of viral RNA template. Thermocycling conditions were 50°C for 10 min, 95°C for 1 min, and 39 cycles of 95°C for 10 s and 65°C for 30 s. Data were analyzed in the CFX Maestro Software (version 2.3) [83]. Thresholds were applied based on the negative controls.

### Evaluation of the frequency of mutant allele over time

The frequency of the mutant allele was calculated as the proportion of total genome copies of virus that were derived from the Tx/12 mutant BC2 virus, calculated as Tx/12 mutant BC2 genome copies divided by the sum of Tx/12 HA WT BC1 and Tx/12 mutant BC2 genome copies, as determined by ddPCR. Genome copies that fell below the limit of detection, as determined by negative control genome copies, were assigned a 0% frequency. The frequency over time was plotted using the tidyverse package version 2.0.0 in RStudio.

### Estimating growth rates of Tx/12 HA F159S BC2 and Tx/12 HA A138S/F159S/N225D BC2 viruses

To infer the variant proportion growth rates, we fit generalized linear mixed model in R version 4.5.1. glmmTMB was used to fit beta-binomial model to barcode counts data. Samples with 0 barcode counts either for the wt or the mutant variant were discarded. The inoculum frequencies at day 0 were used as offset.

We modeled the number of mutant barcode reads *Y_ekt_* as *Y_ekt_* ∼ BetaBinomial(*N_ekt_*, *p_ekt_*, *ϕ*) where *e* is the experiment, *k* is the experimental setting (defined by time and dosage of mAb 2A05 delivery and whether the single or triple mutant virus was used), *N_ekt_* is the total number of mutant and wildtype barcode reads, *p_ekt_* is the expected mutant frequency and *ϕ* is the beta-binomial dispersion parameter. *p_ekt_*was modeled as logit(*p_ekt_*) = logit(*p*_0,*e*_) + (*β_K_* + *b_e_*)*t* where *p*_(0,*e*)_ is the mutant frequency in the inoculum of experiment *e* (0.1 or 0.5), *β_k_* is the fixed-effect slope for experimental setting *k* , and *b_e_* is an experiment-specific random deviation from that slope. The random slopes were assumed to follow *b_e_* ∼ *N*(0, *σ*_*b*_^2^).

### Quantitative analysis of compensatory mutations captured in whole genome sequencing

The proportion of compensatory mutations (*S*) was estimated using the Poisson-based cumulative coverage equation:

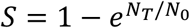

where *N_T_* is the total number of mutants observed and *N_0_* is the total number of unique mutations observed.

### Area plot of amino acid changes in HA over time

A total of 31,880 genomic sequences of the HA gene segment from A/H3N2 were used from GenBank from 2010 through 2020. The sequences were categorized by the amino acids present at position 138, 159, and 225 and plotted across year.

### Phylogenetic tree construction

A total of 381 amino acid sequences of the HA gene segment from A/H3N2 were used from GenBank [36]. The sequences were aligned using MEGA12 version 12.1.2 [84] and the alignment was used to construct the maximum likelihood phylogenetic tree in RAxML-NG version 1.2.1 [85] with 1000 bootstrap replicates (Felsenstein Bootstrap) using the Jones-Taylor-Thornton substitution matrix [86] with a gamma-distributed rate heterogeneity of 4 [87]. The maximum likelihood phylogenetic tree was then visualized using FigTree version 1.4.4 [88].

### Estimation of N_e_

Effective population size was estimated using a previously published method [37]. Simulations were run using a time interval of 3 days, with a minimum read count per barcode of three and minimum frequencies of 10^-5^, 10^-4^, and 10^-3^.

### Wright-Fisher simulations

Wright-Fisher dynamics were simulated at different population sizes and initial starting frequencies using a custom python script, available in our FigShare repository.

### Evaluation of the relationship between N_e_ estimate and viral titer

Estimates of N_e_ and viral titers (determined by plaque assay or ddPCR genome copies) were log transformed and plotted using the tidyverse package version 2.0.0 and ggpubr package version 0.6.0 in RStudio. Linear regression was performed using the ordinary least squares method.

