## Supplement 1 for "Within-host antigenic selection of influenza A virus dominates over stochasticity but is limited by fitness tradeoffs and timing of the immune response"

### Supplemental Table 1: Composition of viral inocula.

<table>
<tr><th>Viruses in inoculum</th><th>Target inoculum composition</th><th>% Tx/12 WT BC1 virus</th><th>% Tx/12 HA F159S BC2 virus</th><th>% Tx/12 HA A138S/ F159S/ N225D BC2 virus</th><th>Barcode distribution[1]</th><th>Figure reference</th><th>Exp. No.</th></tr>
<tr><td rowspan="6">Tx/12 WT BC1 + Tx/12 HA F159S BC2</td><td rowspan="6">90% Tx/12 WT BC1 + 10%Tx/12 HA F159S BC2</td><td>No data</td><td>No data</td><td>-</td><td>No data</td><td>Fig 2<br>Suppl. Fig 3</td><td>1</td></tr>
<tr><td>No data</td><td>No data</td><td>-</td><td>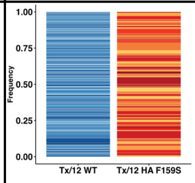
</td><td>Fig 2<br>Suppl. Fig 3</td><td>2</td></tr>
<tr><td>87.5</td><td>12.5</td><td>-</td><td>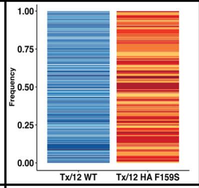
</td><td>Fig 2<br>Suppl. Fig 3</td><td>3</td></tr>
<tr><td>93.8</td><td>6.3</td><td>-</td><td>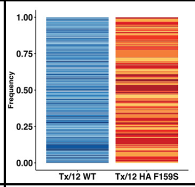
</td><td>Fig 3<br>Suppl. Fig 5</td><td>4</td></tr>
<tr><td>91.0</td><td>9.0</td><td>-</td><td>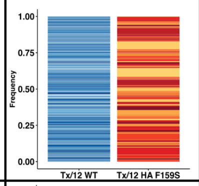
</td><td>Fig 2<br>Suppl. Fig 3</td><td>8</td></tr>
<tr><td>85.0</td><td>15.0</td><td>-</td><td>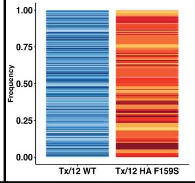
</td><td>Fig 3<br>Suppl. Fig 5</td><td>9</td></tr>
<tr><td>Tx/12 WT BC1 + Tx/12 HA F159S BC2</td><td>50% Tx/12 WT BC1 + 50% Tx/12 HA F159S BC2</td><td>48.8</td><td>51.2</td><td>-</td><td>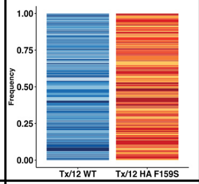
</td><td>Suppl. Fig 2<br>Suppl. Fig 4</td><td>5, 6</td></tr>
<tr><td>Tx/12 WT BC1 + Tx/12 HA A138S/F159 S/N225D BC2</td><td>50% Tx/12 WT BC1 + 50% Tx/12HA A138S/F159 S/N225D BC2</td><td>40.4</td><td>-</td><td>59.6</td><td>No sequence data</td><td>Fig 4</td><td>N/A (did not perform BC sequencing)</td></tr>
<tr><td>Tx/12 WT BC1 + Tx/12 HA A138S/F159 S/N225D BC2</td><td>90% Tx/12 WT BC1 + 10% Tx/12 HA A138S/F159 S/N225D BC2</td><td>85.1</td><td>-</td><td>14.9</td><td>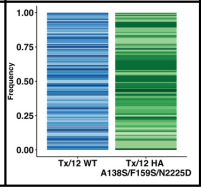
</td><td>Fig 5<br>Suppl. Fig 9</td><td>10, 11</td></tr>
</table>

[1]In stacked barplots, each colored section represents a different barcode, and its height is its relative frequency within the subpopulation.

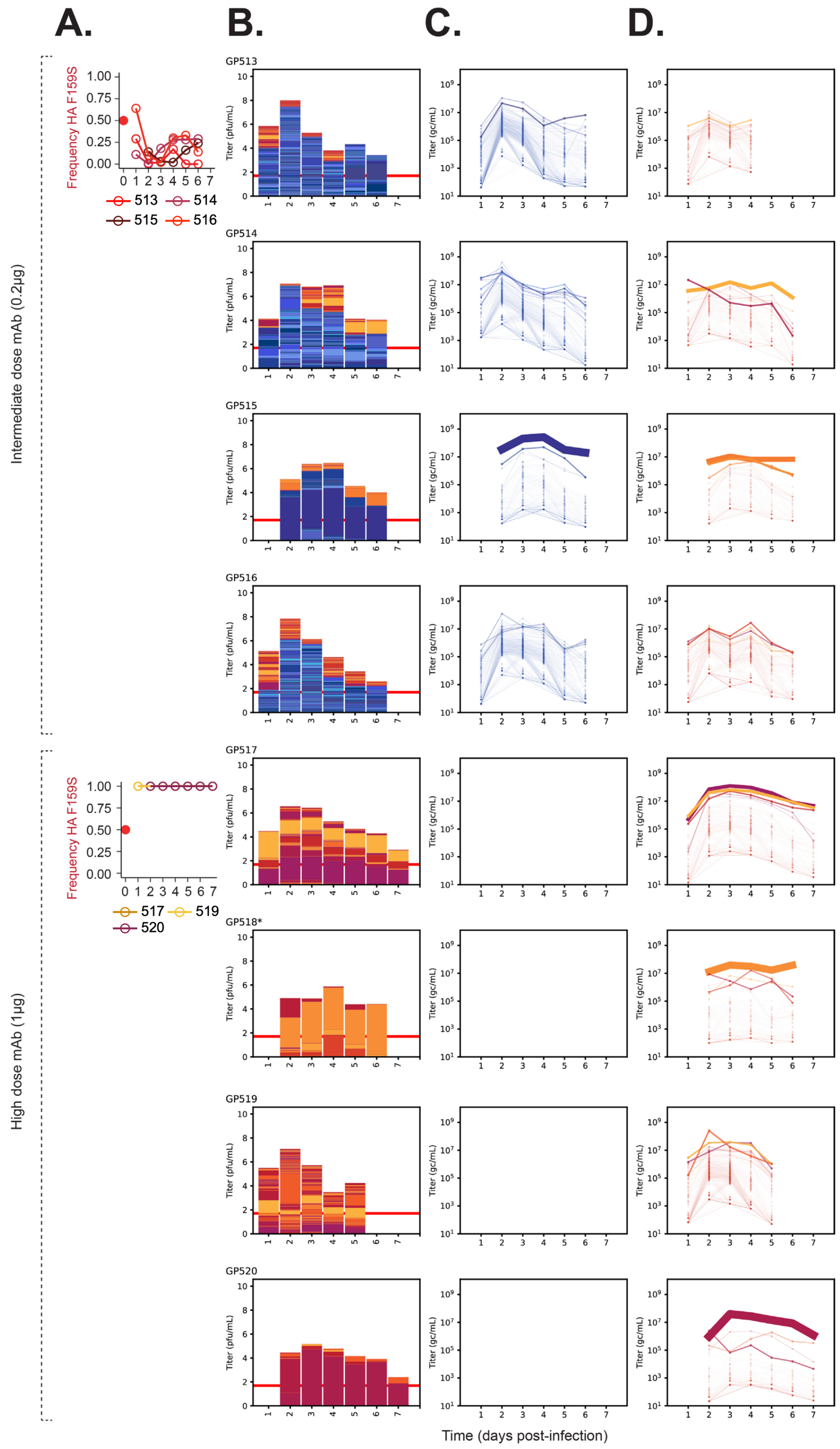

A.
B.
C.
D.
Intermediate dose mAb (0.2μg)
High dose mAb (1μg)
Frequency HA F159S
513
514
515
516
517
519
520
GP513
GP514
GP515
GP516
GP517
GP518*
GP519
GP520
Titer (pfu/mL)
Titer (gc/mL)
Time (days post-infection)

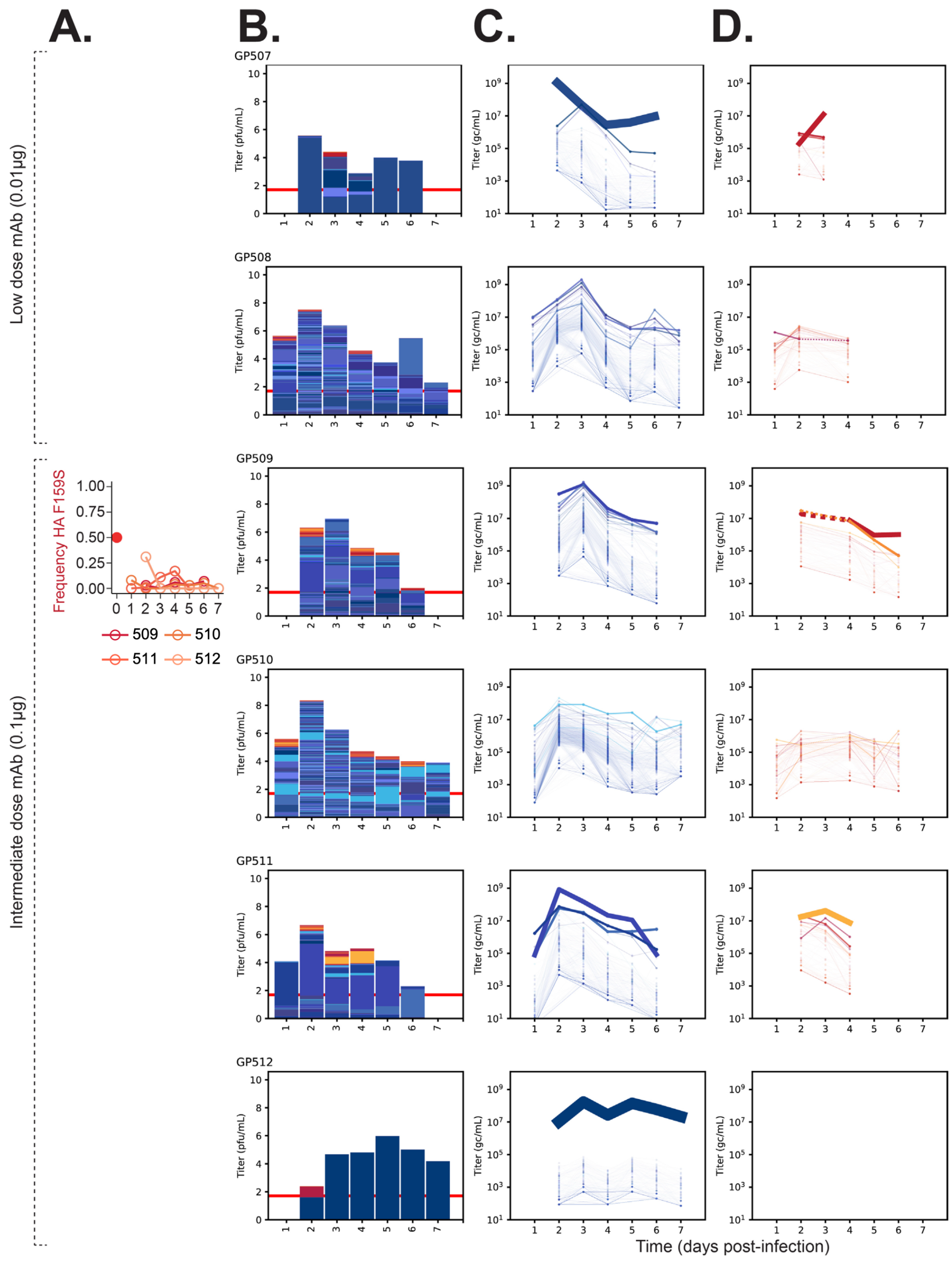

A.
B.
C.
D.
Low dose mAb (0.01μg)
Intermediate dose mAb (0.1μg)
Frequency HA F159S
509
510
511
512
GP507
GP508
GP509
GP510
GP511
GP512
Titer (pfu/mL)
Titer (gc/mL)
Time (days post-infection)

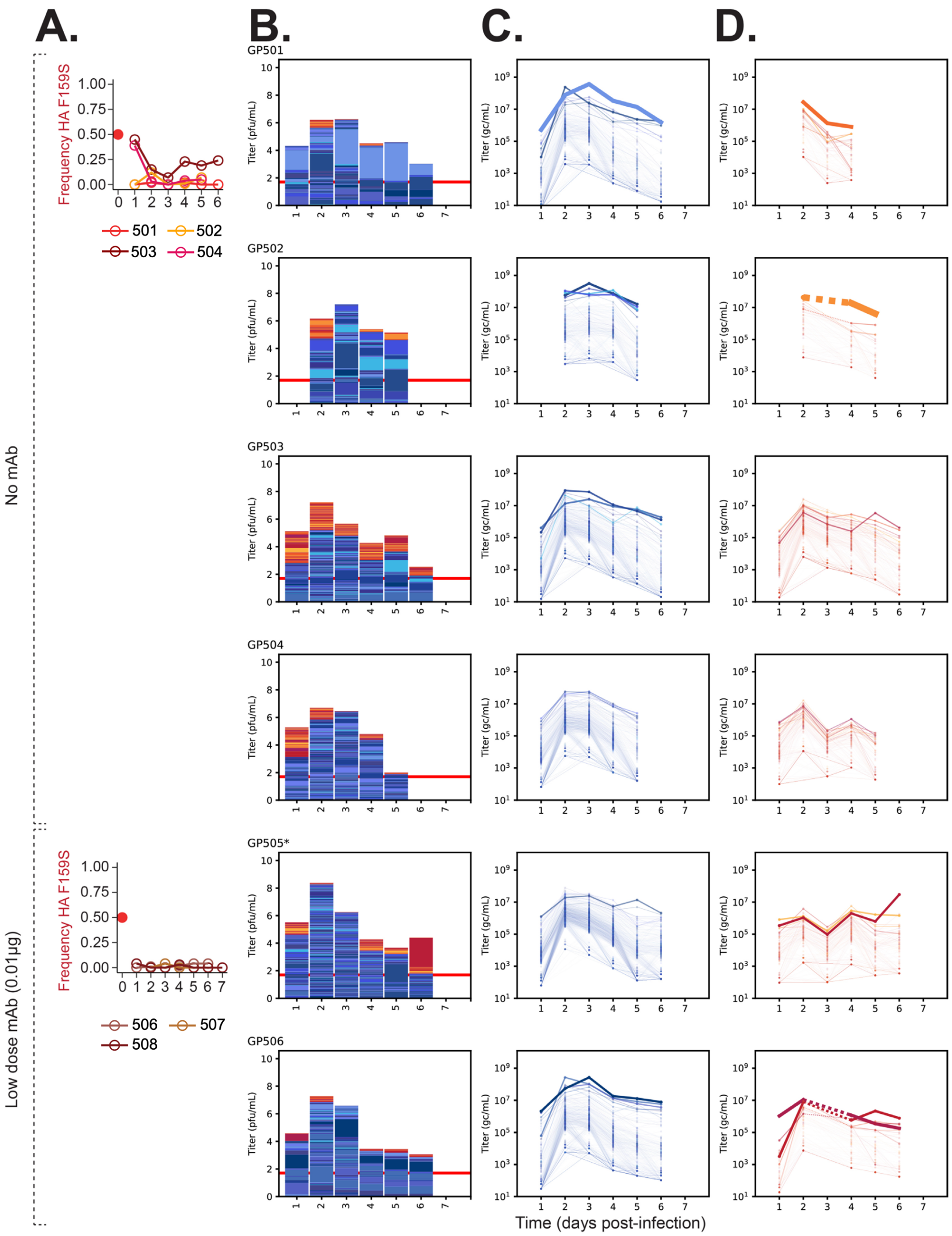

A.
B.
C.
D.
No mAb
Low dose mAb (0.01μg)
Frequency HA F159S
501
502
503
504
506
507
508
GP501
GP502
GP503
GP504
GP505*
GP506
Titer (pfu/mL)
Titer (gc/mL)
Time (days post-infection)

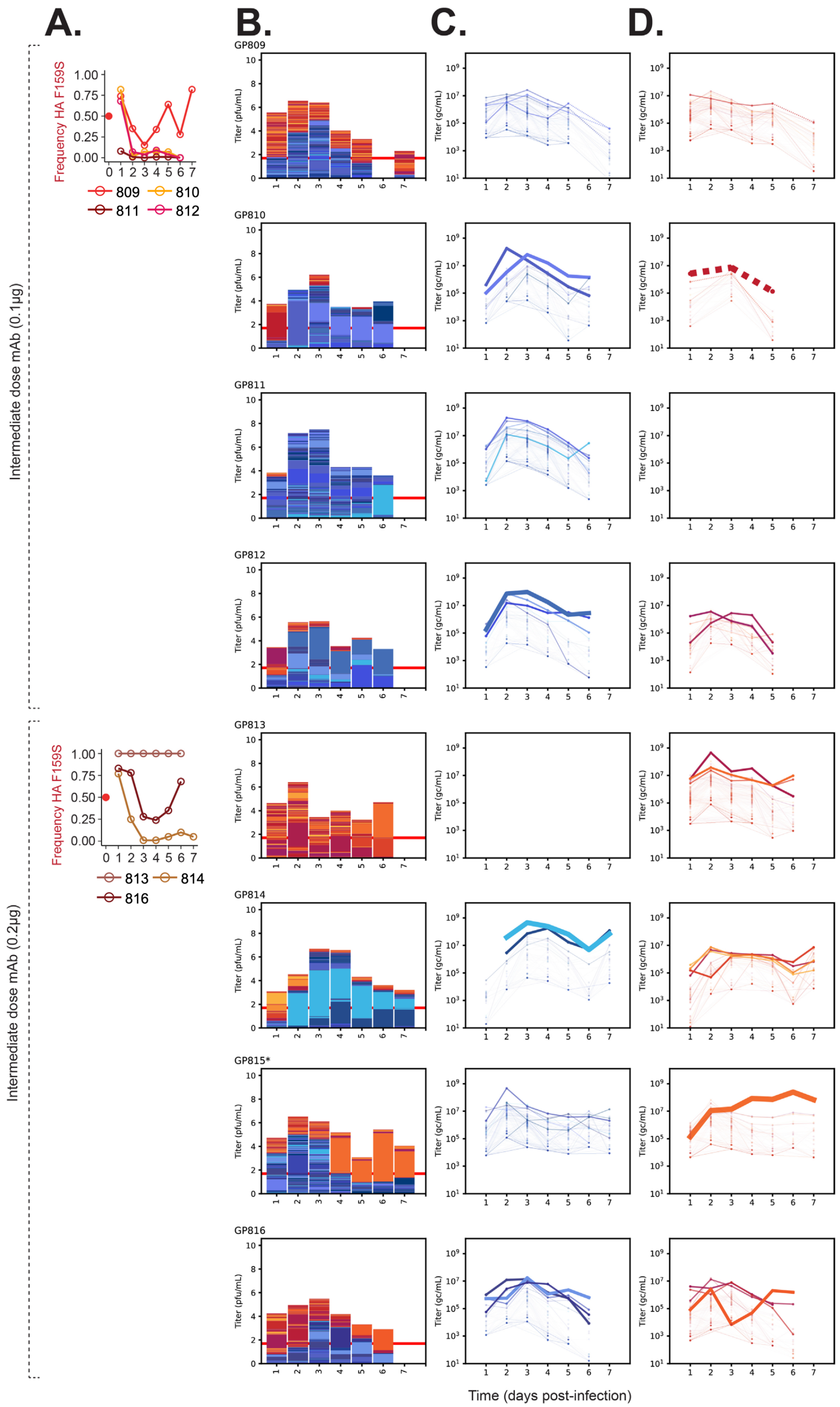

A.
B.
C.
D.
Intermediate dose mAb (0.1µg)
Intermediate dose mAb (0.2µg)
Frequency HA F159S
809
810
811
812
813
814
816
GP809
GP810
GP811
GP812
GP813
GP814
GP815*
GP816
Titer (pfu/mL)
Titer (gc/mL)
Time (days post-infection)

**Supplemental Figure 1: With high initial frequency of the Tx/12 HA F159S BC2 variant, deterministic forces shape overall antigenic variant abundance despite concurrent stochasticity within subpopulations.** Tx/12 HA WT BC1 virus is represented by blue/purple colors and Tx/12 HA F159S BC2 is represented by red/yellow colors. Animals were inoculated with 50% of both viruses. Animals marked with an asterisk had *de novo* mutations that arose and swept with a barcode. **(A)** Frequency of the Tx/12 HA F159S allele over days of infection. Solid dot denotes starting frequency of Tx/12 HA F159S in the inoculum. Each shade of red represents a different animal. Animals in which *de novo* mutations are associated with barcode dynamics are not displayed. Panels **(B)-(D)** contain all animals (the same animal across panels). **(B)** Stacked bar plots where the total height of the column is scaled to viral titer ($log_{10}$ pfu/ml). Each colored section within the column represents a unique barcode, and its height is the relative frequency within the sample. The total height of red/yellow bars and blue/purple bars is scaled to the gc/ml titer of each virus. Red horizontal line represents limit of detection of plaque assay (50 pfu/ml). **(C)** and **(D)** show Tx/12 HA WT and F159S individual barcodes over time, respectively. Each line is a unique barcode with its frequency scaled to the gc/ml titer. Line thickness is scaled to the average frequency of the barcode. Gaps denote results below the limit of detection. Dashed lines connect data separated by single days below the limit of detection.

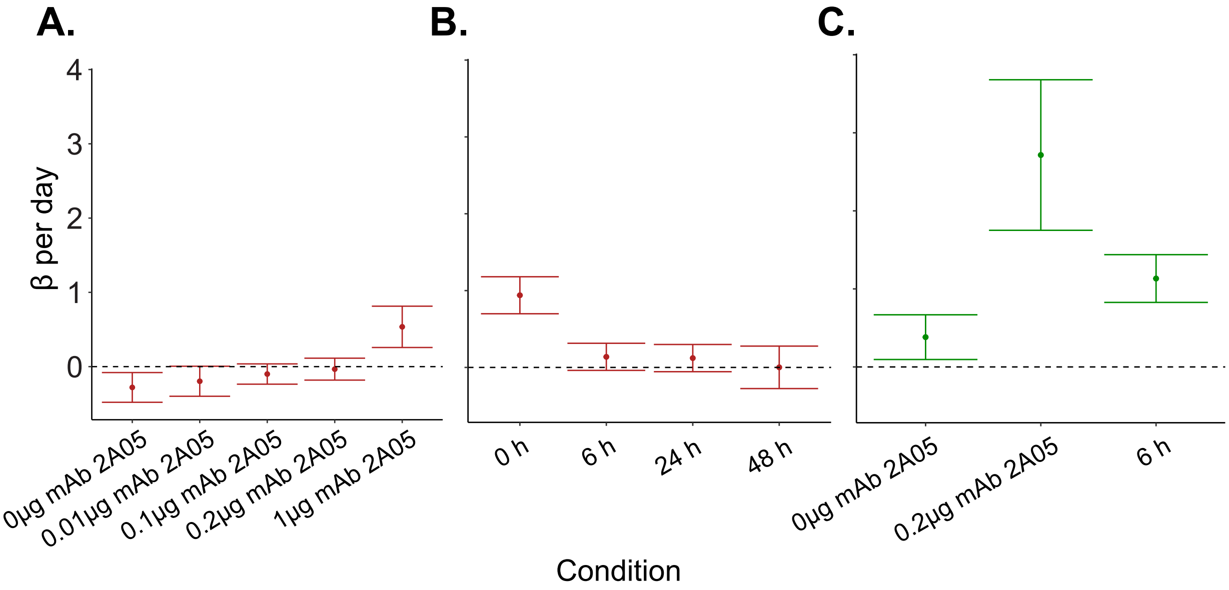


**Supplemental Figure 2: Antigenic variant growth rates are modulated by the strength and timing of immune pressure, and the extent of fitness tradeoffs**. Estimates of β growth rates per day using barcode read counts fit to a beta-binomial model **(A)** at different doses of mAb 2A05, **(B)** given delayed introductions of mAb 2A05, and **(C)** when fitness tradeoffs are eased with the addition of second-site mutations. For panels (A) and (B), data from animals inoculated with 10% or 50% starting frequency of the antigenic variant are combined.

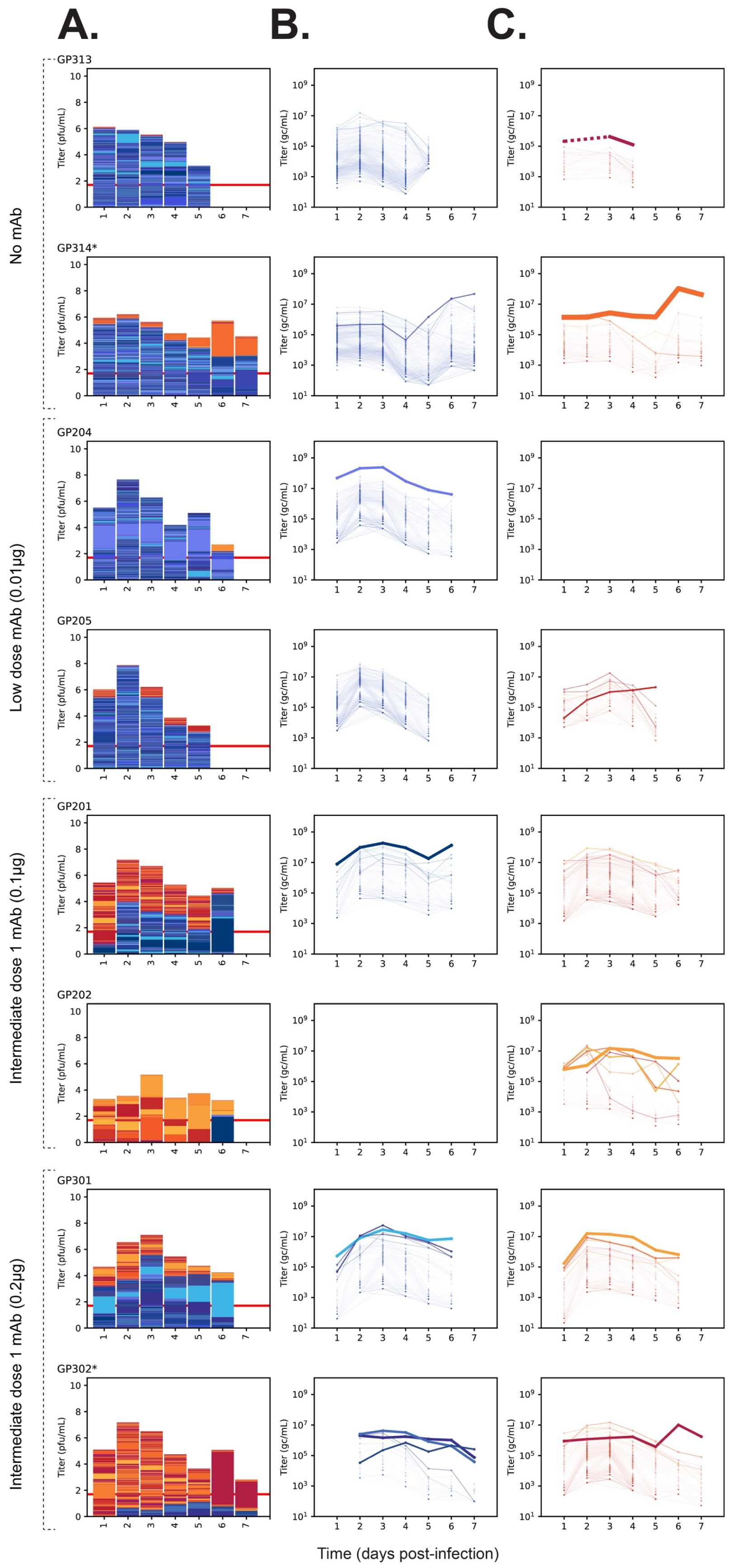

A.
B.
C.
No mAb
Low dose mAb (0.01μg)
Intermediate dose 1 mAb (0.1μg)
Intermediate dose 1 mAb (0.2μg)
GP313
GP314*
GP204
GP205
GP201
GP202
GP301
GP302*
Titer (pfu/mL)
Titer (gc/mL)
Time (days post-infection)

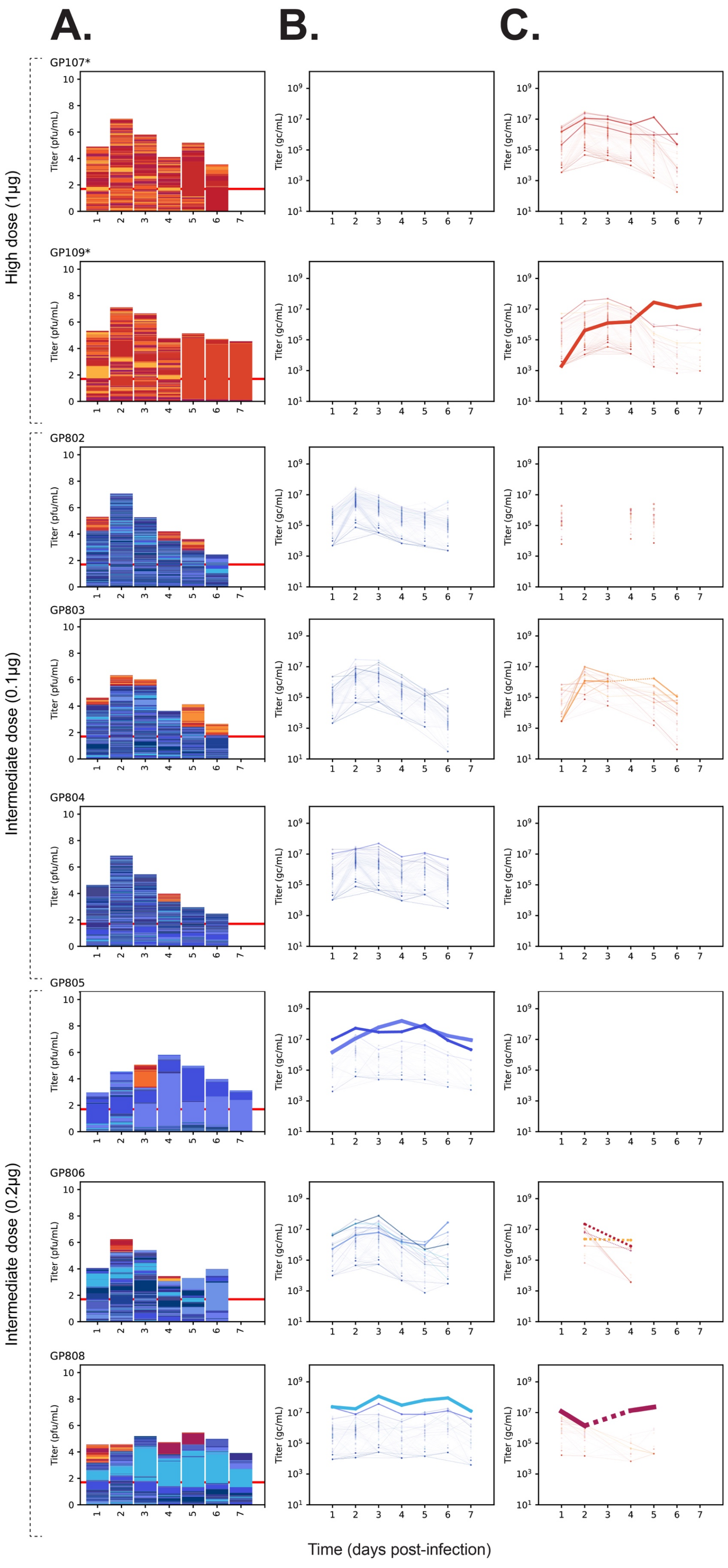
A.
B.
C.
High dose (1μg)
Intermediate dose (0.1μg)
Intermediate dose (0.2μg)
GP107*
GP109*
GP802
GP803
GP804
GP805
GP806
GP808
Titer (pfu/mL)
Titer (gc/mL)
Time (days post-infection)

**Supplemental Figure 3: Deterministic forces shape overall antigenic variant abundance despite some concurrent stochasticity within subpopulations (Related to Figure 2).** Representative animals from these experimental conditions are shown in Figure 2 and remaining animals are shown here. Tx/12 HA WT BC1 virus is represented by blue/purple colors and Tx/12 HA F159S BC2 is represented by red/yellow colors. Animals were inoculated with 90% WT and 10% HA F159S BC viruses. An asterisk denote animals in which *de novo* mutations arose and swept with a barcode. **(A)** Stacked bar plots where the total height of the column is scaled to viral titer ($log_{10}$ pfu/ml). Each colored section within the column represents a unique barcode, and its height is the relative frequency within the sample. The total height of red/yellow bars and blue/purple bars is scaled to the gc/ml titer of each virus. Red horizontal line represents limit of detection of plaque assay (50 pfu/ml). **(B)** and **(C)** show Tx/12 HA WT and F159S individual barcodes over time, respectively. Each line is a unique barcode with its frequency scaled to the gc/ml titer. Line thickness is scaled to the maximum detected frequency of the barcode. Gaps denote results below the limit of detection. Dashed lines connect data separated by single days that fell below the limit of detection.

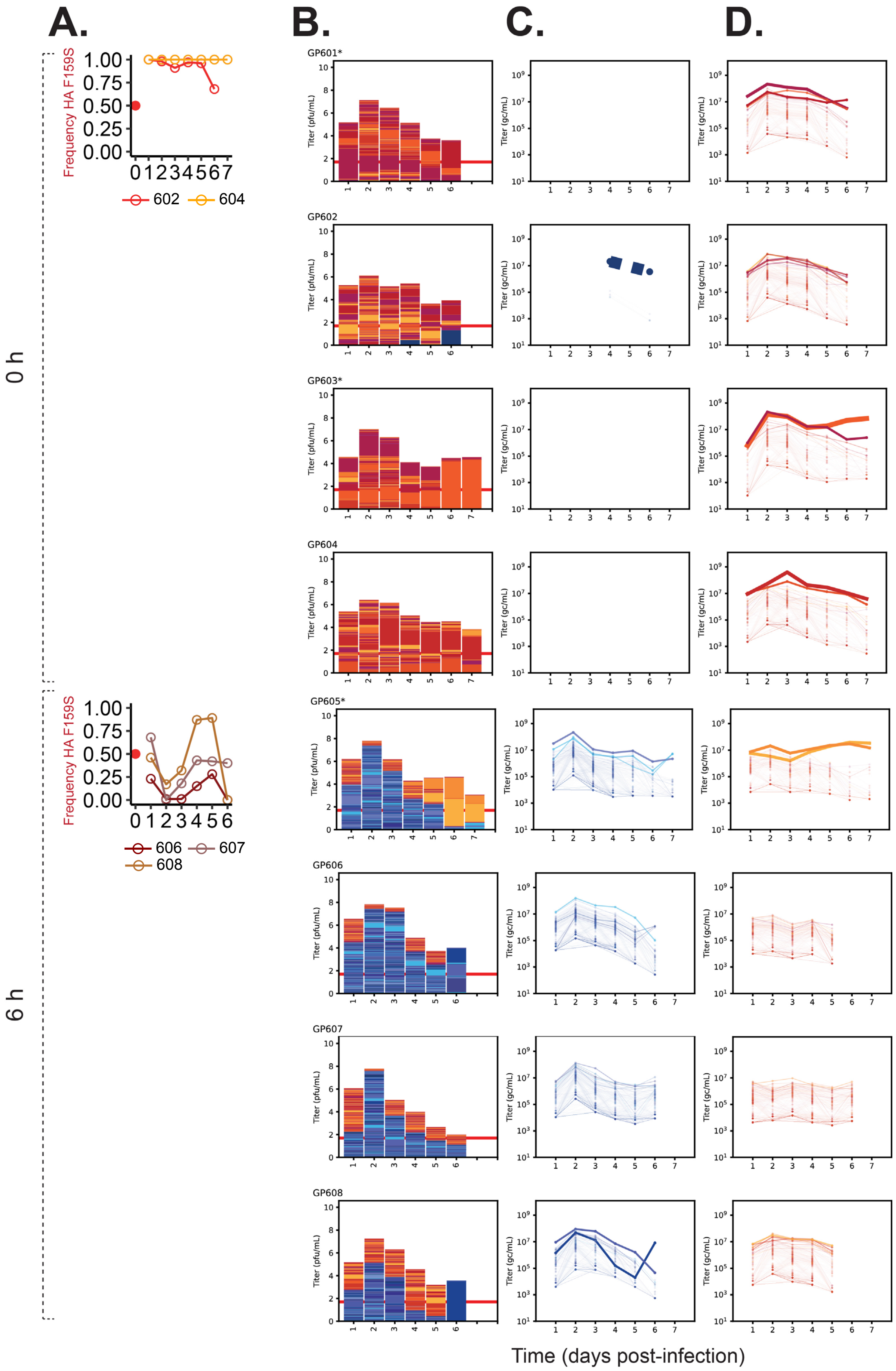
A.
B.
C.
D.
0 h
6 h
Frequency HA F159S
602
604
606
607
608
GP601*
GP602
GP603*
GP604
GP605*
GP606
GP607
GP608
Titer (pfu/mL)
Titer (gc/mL)
Time (days post-infection)

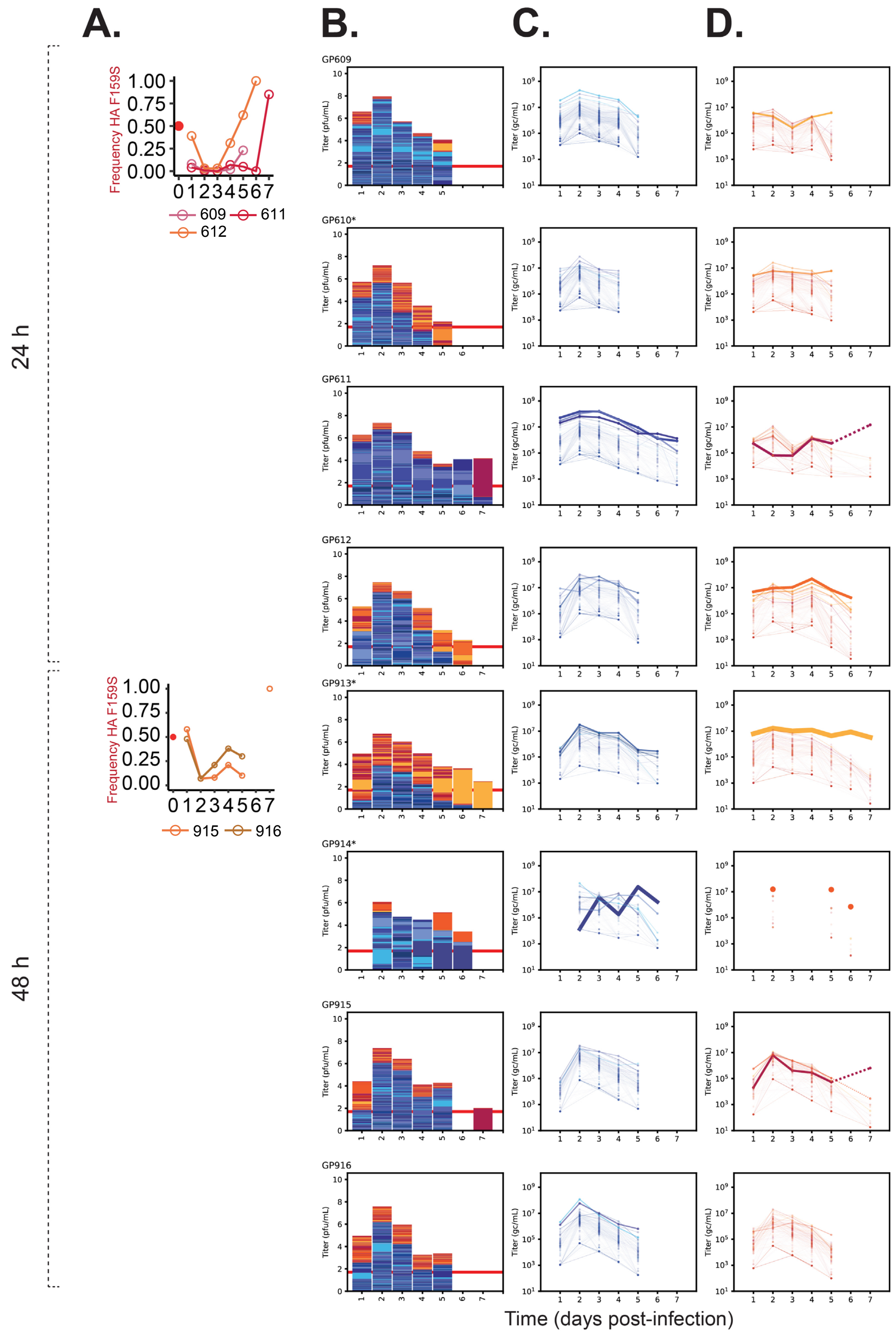

A.
B.
C.
D.
24 h
48 h
Frequency HA F159S
609
611
612
915
916
GP609
GP610*
GP611
GP612
GP913*
GP914*
GP915
GP916
Titer (pfu/mL)
Titer (gc/mL)
Time (days post-infection)

**Supplemental Figure 4: Delay of mAb treatment leads to delayed and inconsistent antigenic selection even at high initial frequency of the Tx/12 HA F159S BC2 variant.** Tx/12 HA WT BC1 virus is represented by blue/purple colors and Tx/12 HA F159S BC2 is represented by red/yellow colors. Animals inoculated with 50% of both viruses. Animals marked with an asterisk had *de novo* mutations that arose and swept with a barcode. **(A)** Frequency of the Tx/12 HA F159S allele over days of infection. Solid dot denotes starting frequency of Tx/12 HA F159S in the inoculum. Each shade of red represents a different animal. Animals in which *de novo* mutations are associated with barcode dynamics are not displayed. **(B)** Stacked bar plots where the total height of the column is scaled to viral titer ($\log_{10}$ pfu/ml). Each colored section within the column represents a unique barcode, and its height is the relative frequency within the sample. The total height of red/yellow bars and blue/purple bars is scaled to the gc/ml titer of each virus. Red horizontal line represents limit of detection of plaque assay (50 pfu/ml). **(C)** and **(D)** show Tx/12 HA WT and F159S individual barcodes over time, respectively. Each line is a unique barcode with its frequency scaled to the gc/ml titer. Line thickness is scaled to the average frequency of the barcode. Gaps denote results below the limit of detection.

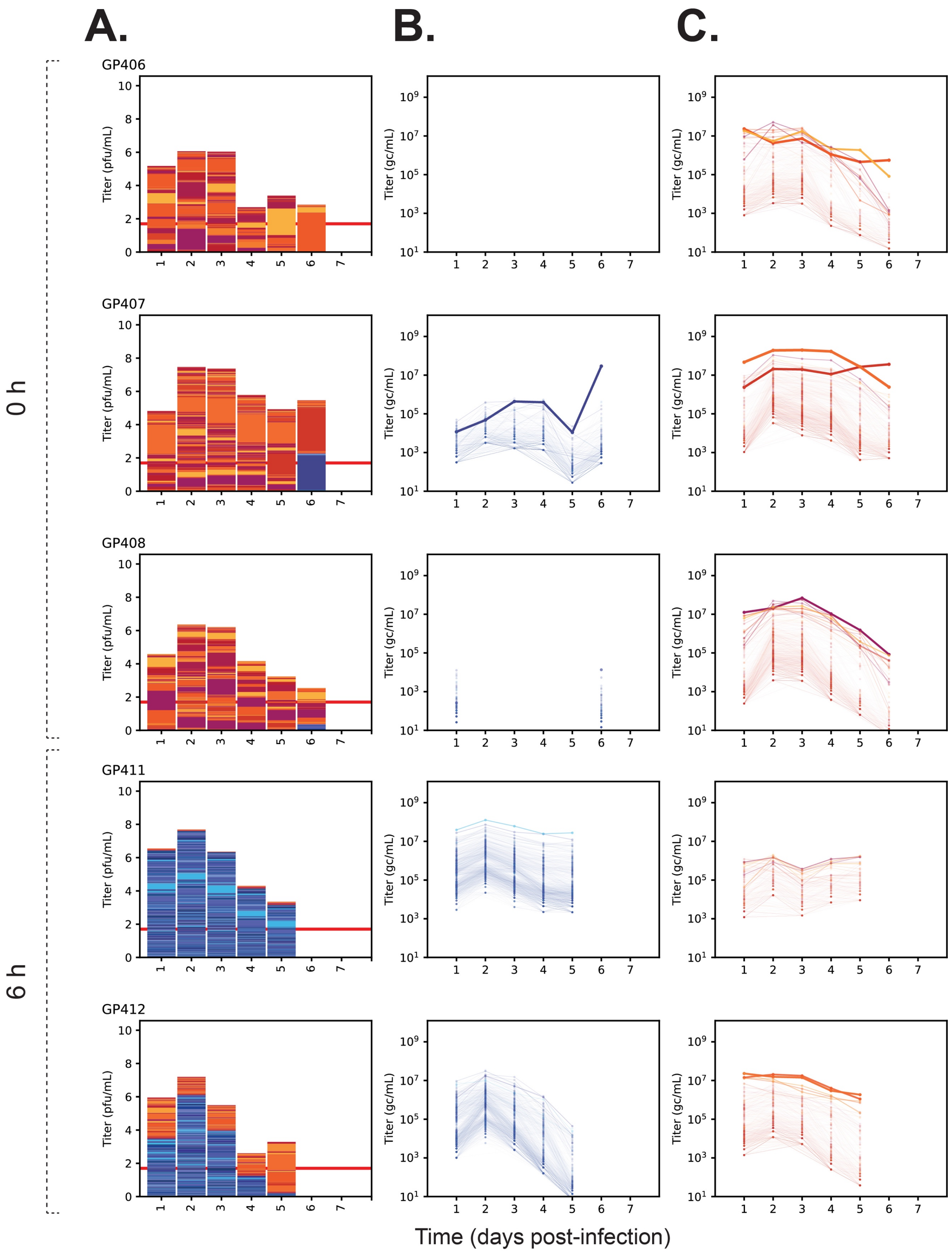
A.
B.
C.
0 h
6 h
GP406
GP407
GP408
GP411
GP412
Titer (pfu/mL)
Titer (gc/mL)
Time (days post-infection)

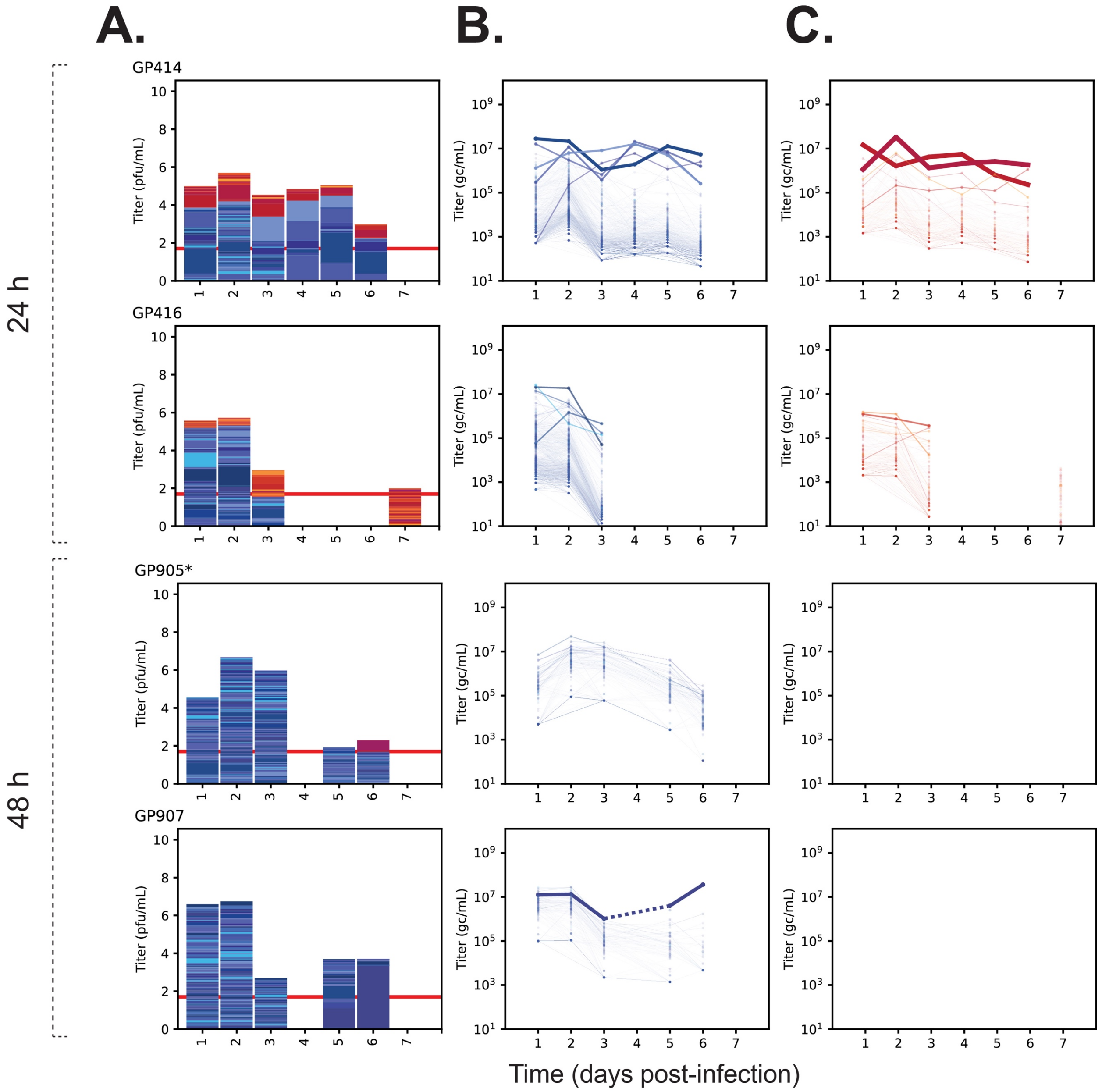


**Supplemental Figure 5: Delay of mAb treatment leads to delayed and inconsistent antigenic selection (Related to Figure 3).** Representative animals from these experimental conditions are shown in Figure 2 and remaining animals are shown here. Tx/12 HA WT BC1 virus is represented by blue/purple colors and Tx/12 HA F159S BC2 is represented by red/yellow colors. Animals were inoculated with 90% WT and 10% HA F159S BC viruses.An asterisk denotes animals in which *de novo* mutation(s) arose and swept with a barcode. **(A)** Stacked bar plots where the total height of the column is scaled to viral titer ($log_{10}$ pfu/ml). Each colored section within the column represents a unique barcode, and its height is the relative frequency within the sample. The total height of red/yellow bars and blue/purple bars is scaled to the gc/ml titer of each virus. Red horizontal line represents limit of detection of plaque assay (50 pfu/ml). **(B)** and **(C)** show individual Tx/12 HA WT and F159S barcodes over time, respectively. Each line is a unique barcode with its frequency scaled to the gc/ml titer. Line thickness is scaled to the maximum detected frequency of the barcode. Gaps denote results below the limit of detection. Dashed lines connect data separated by single days that fell below the limit of detection.

#### Supplemental Table 2: *De novo* mutations linked to a barcode arise in a subset of animals across conditions largely in the Tx/12 HA F159S background

| GP | Inoculum | mAb condition | BC dynamics | *De novo* mutations linked to BC (H3 numbering)[1] | % WGS iSNV | BC number | % BC | Conclusion |
|---|---|---|---|---|---|---|---|---|
| 313 | 10-1 | None | Expected | N/A | N/A | N/A | N/A | BCs neutral |
| 314 | 10-1 | None | Outgrowth of HA F159S BC | I140K | Day 2: not detected<br>Day 4: 10%<br>Day 6: 48.9% | HABC2-595 | Day 2: 18% of S159 subpopulation, S159 2% of total virus population<br>Day 4: 74% of S159 subpopulation, S159 10% of total virus population<br>Day 6: 91% of S159 subpopulation, S159 46% of total virus population | BC not neutral |
| 205 | 10-1 | Low dose | Expected | N/A | N/A | N/A | N/A | BCs neutral |
| 203 | 10-1 | Intermediate dose 1 | Expected | N/A | N/A | N/A | N/A | BCs neutral |
| 302 | 10-1 | Intermediate dose 2 | Outgrowth HA F159S BC | I226N | Day 2: not detected<br>Day 4: not detected<br>Day 6: 83% | HABC2-544 | Day 2: 1%<br>Day 4: 5.6%<br>Day 6: 93.4% | BC not neutral |
| 303 | 10-1 | Intermediate dose 2 | Expected | N/A | N/A | N/A | N/A | BCs neutral |
| 107 | 10-1 | High dose | Largely expected, some potential outgrowth of HA F159S BC | V186I | Day 2: not detected<br>Day 4: not detected<br>Day 6: 100% | HABC2-27 | Day 2: 0.7%<br>Day 4: 0.8%<br>Day 6: 51% | BC not neutral |
| 109 | 10-1 | High dose | Outgrowth of HA F159S BC | H156R | Day 2: not detected<br>Day 4: not detected<br>Day 6: 89% | HABC2-145 | Day 2: 0.2%<br>Day 4: 0.9%<br>Day 6: 88% | BC not neutral |
| 407 | 10-1 | 0 h instillation | Expected | N/A | N/A | N/A | N/A | BCs neutral |
| 405 | 10-1 | 0 h instillation | Outgrowth of HA F159S BC | N/A | N/A | HABC2-336 | Day 2: 34.9%<br>Day 4: 62.6%<br>Day 6: 78% | BCs neutral |
| 410 | 10-1 | 6 h instillation | Expected | N/A | N/A | N/A | N/A | BCs neutral |
| 409 | 10-1 | 6 h instillation | Outgrowth of WT BC | N/A | N/A | HABC1-934 | Day 1: 10%<br>Day 3: 74.5%<br>Day 5: 73% | BCs neutral |
| 413 | 10-1 | 24 h instillation | Expected | N/A | N/A | N/A | N/A | BCs neutral |
| 503_ seq 1 | 1-1 | None | Expected | N/A | N/A | N/A | N/A | BCs neutral; low coverage |
| 503_ seq 2 | 1-1 | None | Expected | N/A | N/A | N/A | N/A | BCs neutral; repeat sequencing |

| | | | | | | | | |
|---|---|---|---|---|---|---|---|---|
| 505 | 1-1 | Low dose | Outgrowth of HA F159S BC | H156R | Day 2: not detected<br>Day 4: not detected<br>Day 6: 73% | HABC2-1008 | Day 2: 3.3%<br>Day 4: 4.1%<br>Day 6: 81% | BC not neutral |
| 506 | 1-1 | Low dose | Expected | N/A | N/A | N/A | N/A | BCs neutral |
| 507_ seq 1 | 1-1 | Low dose | Outgrowth of WT BC | N/A | N/A | HABC1-74 | Day 2: 97%,<br>Day 4: 44.6%<br>Day 6: 98.6% | BCs neutral; low coverage |
| 507_ seq 2 | 1-1 | Low dose | Outgrowth of WT BC | N/A | N/A | HABC1-74 | Day 2: 97%,<br>Day 4: 44.6%,<br>Day 6: 98.6% | BCs neutral; repeat sequencing |
| 518_ seq 1 | 1-1 | High dose | Outgrowth of HA F159S BC | HA2 D157N | Day 2: not detected<br>Day 4: ~44%<br>Day 6: not detected | HABC2-927 | Day 2: 50.9%<br>Day 4: 59.7%<br>Day 6: 94.5% | BC not neutral; low coverage |
| 518_ seq 2 | 1-1 | High dose | Outgrowth HA F159S BC | HA2 D157N | Day 2: 43%<br>Day 4: 40%<br>Day 6: 94% | HABC2-927 | Day 2: 50.9%<br>Day 4: 59.7%<br>Day 6: 94.5% | BC not neutral; repeated sequencing |
| 519 | 1-1 | High dose | Expected | N/A | N/A | N/A | N/A | BCs neutral |
| 520_ seq 1 | 1-1 | High dose | Outgrowth of HA F159S BC | N/A | N/A | HABC2-1 | Day 2: 24.5%<br>Day 4: 86.2%<br>Day 6: 93.1% | BCs neutral; low coverage |
| 520_ seq 2 | 1-1 | High dose | Outgrowth of HA F159S BC | N/A | N/A | HABC2-1 | Day 2: 24.5%<br>Day 4: 86.2%<br>Day 6: 93.1% | BCs neutral; repeat sequencing |
| 601 | 1-1 | 0 h instillation | Largely expected, some potential outgrowth of HA F159S BC | T248I | Day 2: not detected<br>Day 4: not detected<br>Day 6: ~80% | HABC2-560 | Day 2: 4.9%<br>Day 4: 4.1%<br>Day 6: ~62.7% | BCs not neutral |
| 603 | 1-1 | 0 h instillation | Outgrowth of HA F159S BC | N246H | Day 2: not detected<br>Day 4: not detected<br>Day 6: 99% | HABC2-356 | Day 2: 16.6%<br>Day 4: 22%<br>Day 6: 91.4% | BCs not neutral |
| 605 | 1-1 | 6 h instillation | Outgrowth of HA F159S BC | H156R, H156Q | Day 2: not detected<br>Day 4: no data<br>Day 6: 56%, 40% | HABC2-439, HABC2-927 | Day 2: 2.1%, 12.3%<br>Day 4: 13.6%, 21.3%<br>Day 6: 55.3%, 42% | BCs not neutral |
| 606 | 1-1 | 6 h instillation | Outgrowth WT BC on last day | N/A | N/A | HABC1-937, HABC1-44, HABC1-559 | Day 2: 0.27%, 0.44%, 0.21%<br>Day 4: 0.34%, 0.93%, 0.23%<br>Day 6: 29.7%, 29.4%, 24.2% | BCs neutral |
| 608 | 1-1 | 6 h instillation | Largely expected, some outgrowth of WT BC on last day | N/A | N/A | HABC1-687 | Day 2: 3.5%<br>Day 4: 0.38%<br>Day 6: 98.1% | BCs neutral |
| 609 | 1-1 | 24 h instillation | Expected | N/A | N/A | N/A | N/A | BCs neutral |
| 610 | 1-1 | 24 h instillation | Outgrowth of HA F159S BC | NA K308N | Day 1: not detected<br>Day 3: not detected | HABC2-565 | Day 1: 6.6%<br>Day 3: 3.3%<br>Day 5: 46.4% | BC may be linked to NA mutation |

| | | | | | | | | |
|---|---|---|---|---|---|---|---|---|
| | | | | | Day 5: ~67% | | | |
| 611 | 1-1 | 24 h instillation | Outgrowth HA F159S BC | N/A | N/A | HABC2-569 | Day 2: 0.046%<br>Day 4: 5.0%<br>Day 6: 25.9%<br>Day 7: 98.2% of total | BCs neutral |
| 509 | 1-1 | Intermediate dose 1 | Expected | N/A | N/A | N/A | N/A | BCs neutral |
| 511 | 1-1 | Intermediate dose 1 | Outgrowth WT BC | N/A | N/A | HABC1-569 | Day 1: 3.5%<br>Day 3: 47.4%<br>Day 5: 68.7% | BCs neutral |
| 512 | 1-1 | Intermediate dose 1 | Outgrowth WT BC | N/A | N/A | HABC1-825 | Day 2, 4, 6: >97% | BCs neutral |
| 514 | 1-1 | Intermediate dose 2 | Expected | N/A | N/A | N/A | N/A | BCs neutral |
| 515 | 1-1 | Intermediate dose 2 | Outgrowth WT BC | N/A | N/A | HABC1-771 | Day 2: 85.4%<br>Day 4: 66.1%<br>Day 6: 94.1% | BCs neutral |
| 702 | WT | None | Expected | N/A | N/A | N/A | N/A | BCs neutral |
| 705 | HA F159S | None | Expected | N/A | N/A | N/A | N/A | BCs neutral |
| 708 | HA F159S | None | Outgrowth HA F159S BC | I226N | Day 2: not detected<br>Day 4: 10%<br>Day 6: 100% | HABC2-751 | Day 2: 0.8%<br>Day 4: 3.3%<br>Day 6: 98.4% | BC not neutral |
| 815 | 1-1 | Intermediate dose 2 | Outgrowth HA F159S BC | H156R | Day 2: not detected<br>Day 4: 71%<br>Day 6: 83% | HABC2-595 | Day 2: 2.3%<br>Day 4: 80.5%<br>Day 6: 92.1% | BC not neutral |
| 801 | 10-1 | Intermediate dose 1 | Expected | N/A | N/A | N/A | N/A | BCs neutral; low coverage |
| 801_seq2 | 10-1 | Intermediate dose 1 | Expected | N/A | N/A | N/A | N/A | BCs neutral; repeat sequencing |
| 805 | 10-1 | Intermediate dose 2 | Outgrowth of WT BC | N/A | N/A | HABC1-651, HABC1-501 | Day 1: 50.5%, 7.7%<br>Day 3: 25.9%, 52.6%<br>Day 5: 48.7%, 30.7% | BCs neutral |
| 806 | 10-1 | Intermediate dose 2 | Outgrowth of WT BC | N/A | N/A | HABC1-619 | Day 2: 1.1%<br>Day 4: 4.5%<br>Day 6: 52.7% | BCs neutral |
| 807 | 10-1 | Intermediate dose 2 | Expected | N/A | N/A | N/A | N/A | BCs neutral |
| 809 | 1-1 | Intermediate dose 1 | Expected | N/A | N/A | N/A | N/A | BCs neutral |
| 810 | 1-1 | Intermediate dose 1 | Outgrowth of WT BC | N/A | N/A | HABC1-359 | Day 1: 12.9%<br>Day 3: 28.5%<br>Day 5: 60.6% | BCs neutral |
| 811 | 1-1 | Intermediate dose 1 | Outgrowth of WT BC | N/A | N/A | HABC1-497 | Day 2: 1.0%<br>Day 4: 0.74%<br>Day 6: 70.4% | BCs neutral |
| 812 | 1-1 | Intermediate dose 1 | Outgrowth of WT BC | N/A | N/A | HABC1-459 | Day 1: 4.5%<br>Day 3: 5.2%<br>Day 5: 45.1% | BCs neutral |
| 813 | 1-1 | Intermediate dose 2 | Outgrowth HA F159S BC | N/A | N/A | HABC2-576, HABC2-145 | Day 2: 2.6%, 1.6%<br>Day 4: 4.6%, 4.5%<br>Day 6: 60.5%, 32% | BCs neutral |
| 814 | 1-1 | Intermediate dose 2 | Outgrowth of WT BC | N/A | N/A | HABC1-173 | Day 2: 6.4%<br>Day 4: 29.3%<br>Day 6: 49.4% | BCs neutral |
| 816 | 1-1 | Intermediate dose 2 | Outgrowth HA F159S BC | N/A | N/A | HABC2-836 | Day 2: 4.1%<br>Day 4: 1.2%<br>Day 6: 87.3% | BCs neutral |

| | | | | | | | | |
|---|---|---|---|---|---|---|---|---|
| 901 | 10-1 | 0 h | Outgrowth of HA F159S BC (2 barcodes) | N246S T248R | Day 1: not detected Day 3: not detected Day 5: 39.3%, 41.8% | HABC2-824, HABC2-598 | Day 1: 0.15%, 2.1% Day 3: 1.9%, 2.23% Day 5: 29.8%, 29.6% | BCs not neutral |
| 902 | 10-1 | 0 h | Outgrowth of HA F159S | H156R | Day 1: not detected Day 3: not detected Day 5: 8.3% Day 7: 80.3% | HABC2-374 | Day 1: 1.2% Day 3: 0.29% Day 5: 6.13% Day 7: 97.2% | BC not neutral |
| 903 | 10-1 | 0 h | Expected | H156Q V186I | Day 2: not detected Day 4: not detected Day 6: 15.2%, 23.6% | Top BCs: HABC2-645 HABC2-595 HABC2-576 | Day 2: 0.5%, 1.5%, 1.7%<br><br>Day 4: 0.4%, 2.7%, 4.0%<br><br>Day 6: 14.5%, 11.2%, 10.0% | BCs not neutral |
| 904 | 10-1 | 0 h | Outgrowth of HA F159S | N/A | N/A | HABC2-598, HABC2-540 | Day 2: 1.4%, 0% Day 4: 0.56%, 0.23% Day 6: 50.8%, 26.1% | BCs neutral |
| 905 | 10-1 | 48 h | Outgrowth of HA F159S | R269G | Day 2: not detected Day 4: not detected Day 6: 12.3% | HABC2-721 | Day 2: not detected Day 4: not detected Day 6: 83.2% of S159 subpopulation, S159 27% of total virus population | BC not neutral |
| 906 | 10-1 | 48 h | Outgrowth of WT | N/A | N/A | HABC1-25, HABC1-433, HABC1-744, HABC1-1002 | Day 2: 2.3%, 0.8%, 0.8%, 1.6% Day 4: 2.8%, 3.3%, 0.44%, 1.1% Day 6: 20.3%, 20.3%, 15.2%, 9.8% | BCs neutral |
| 907 | 10-1 | 48 h | Outgrowth of WT | N/A | N/A | HABC1-44 | Day 1: 1.5% Day 3: 4.3% Day 5: 27.7% Day 6: 90.7% | BC neutral |
| 909 | 1-1 | 0 h | Expected | N/A | N/A | N/A | N/A | BC neutral |
| 912 | 1-1 | 0 h | Outgrowth of HA F159S | N/A | N/A | HABC2-191 | Day 2: 19.1% Day 4: 11.5% Day 6: 83.3% | BC neutral |
| 913 | 1-1 | 48 h | Outgrowth of HA F159S | H156R | Day 2: not detected Day 4: not detected Day 6: 96.3% | HABC2-439 | Day 2: 6.2% Day 4: 16.3% Day 6: 92.5% | BC not neutral |
| 914 | 1-1 | 48 h | Outgrowth of HA F159S | T167A | Day 2: not detected Day 4: not detected Day 6: 13.8% | HABC2-256 | Day 2: 51.2% of S159 subpopulation, S159 8% of total virus population | BC not neutral |

| | | | | | | | | |
|---|---|---|---|---|---|---|---|---|
| | | | | | | | Day 4: not detected, S159 5% of total virus population<br>Day 6: 98.8% of S159 subpopulation, S159 29% of total virus population | |
| 915 | 1-1 | 48 h | Outgrowth of HA F159S | N/A | N/A | HABC2-57 | Day 1: 1.2%<br>Day 3: 1.0%<br>Day 5: 4.3%<br>Day 7: 98.2% | BC neutral |
| 916 | 1-1 | 48 h | Expected | N/A | N/A | N/A | N/A | BCs neutral |
| 1001 | 10-1, triple mutant | No mAb | Expected | N/A | N/A | N/A | N/A | BCs neutral |
| 1003 | 10-1, triple mutant | No mAb | Outgrowth of HA A138S/F159S/N225D | H156Q | H156Q:<br>Day 2: not detected<br>Day 4: not detected<br>Day 6: 63.3%<br>Day 7: 0%<br><br>A138S:<br>Day 2: 32%<br>Day 4: 27%<br>Day 6: 0%<br>Day 7: 94% | HABC2-618<br>HABC2-610 | HABC2-618:<br>Day 2: 0%<br>Day 4: 2.2%<br>Day 6: 70.9% of triple mutant subpopulation, triple mutant 17% of total virus population<br>Day 7: 0.1%<br><br>HABC2-610:<br>Day 2: 5.9%<br>Day 4: 7.7%<br>Day 6: 13.2%<br>Day 7: 97.1% | BCs may not be neutral |
| 1007 | 10-1, triple mutant | Intermediate dose 2 | Expected | N/A | N/A | N/A | N/A | BCs neutral |
| 1008 | 10-1, triple mutant | Intermediate dose 2 | Outgrowth of WT | N/A | N/A | HABC1-28, HABC1-684, HABC1-32, HABC1-48, HABC1-668 | Day 1: 68.9%, 0%, 0%, 0%, 0%<br>Day 2: 34.1%, 16.2%, 15.6%, 12.7%, 10.7%<br>Day 3: not detected<br>Day 5: not detected | BCs neutral |
| 1101 | 10-1, triple mutant | 0 h | Expected | N/A | N/A | N/A | N/A | BCs neutral |
| 1103 | 10-1, triple mutant | 0 h | Outgrowth of HA triple mutant barcodes | N/A | N/A | N/A | N/A | BCs neutral |
| 1105 | 10-1, triple mutant | 6 h | Outgrowth of HA triple mutant barcodes | N/A | N/A | N/A | N/A | BCs neutral |
| 1106 | 10-1, triple mutant | 6 h | Outgrowth of HA triple mutant barcodes | N/A | N/A | N/A | N/A | BCs neutral |
| 1107 | 10-1, triple mutant | 6 h | Some outgrowth of HA triple mutant barcodes | N/A | N/A | N/A | N/A | BCs neutral |

| | | | | | | | | |
|---|---|---|---|---|---|---|---|---|
| 1108 | 10-1, triple mutant | 6 h | Expected | N/A | N/A | N/A | N/A | BCs neutral |

[1]Defined as *de novo* mutations that have a similar frequency trajectory to a given BC over time

**SEE SEPARATE PDF FILE**

**Supplemental Figure 6: Whole genome sequencing reveals *de novo* mutations that arise and increase in frequency during infection in a subset of animals.** Whole genome sequencing of guinea pig nasal wash samples. Categorized iSNVs (S=synonymous, NS=nonsynonymous, na=UTR) are plotted at their frequency on the left axis and sequencing coverage (gray line) is plotted on the right axis. Segments are identified above each trace.

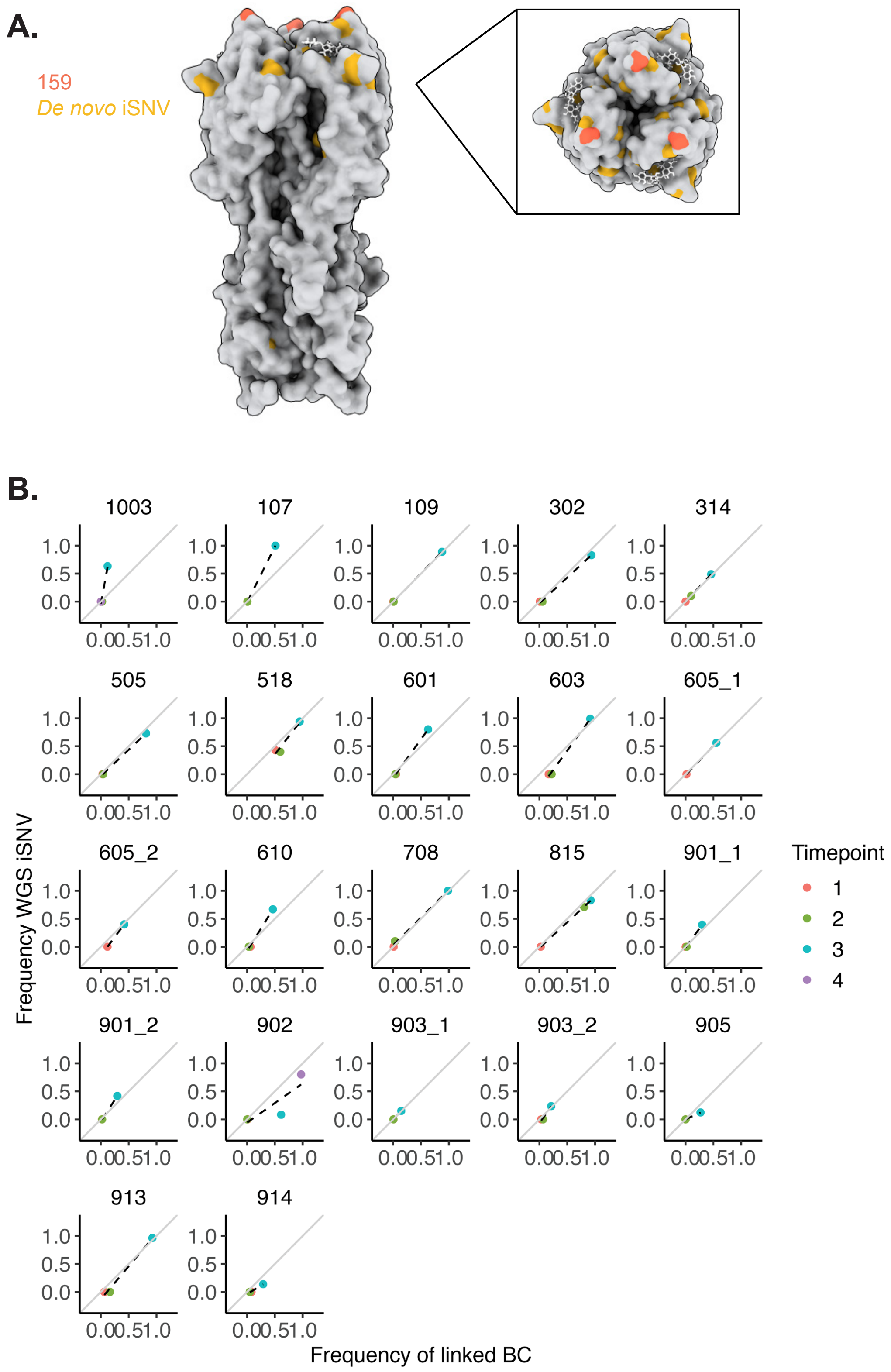


**Supplemental Figure 7: In the HA F159S background, *de novo* mutations clustered around the HA receptor binding site show frequency dynamics similar to those of a given barcode. (A)** Labeled HA trimer at identified *de novo* mutation positions (A/X31,

PDB: 1HGG). Red marks indicate residue 159, yellow marks show residues of identified *de novo* mutations. **(B)** Frequency plots of *de novo* iSNVs and linked barcode motif colored by time sampled. Gray line shows x=y and dashed line is a linear regression fit to the data. Guinea pig number above each trace. Two entries per animal indicate two different barcodes linked to an iSNV.

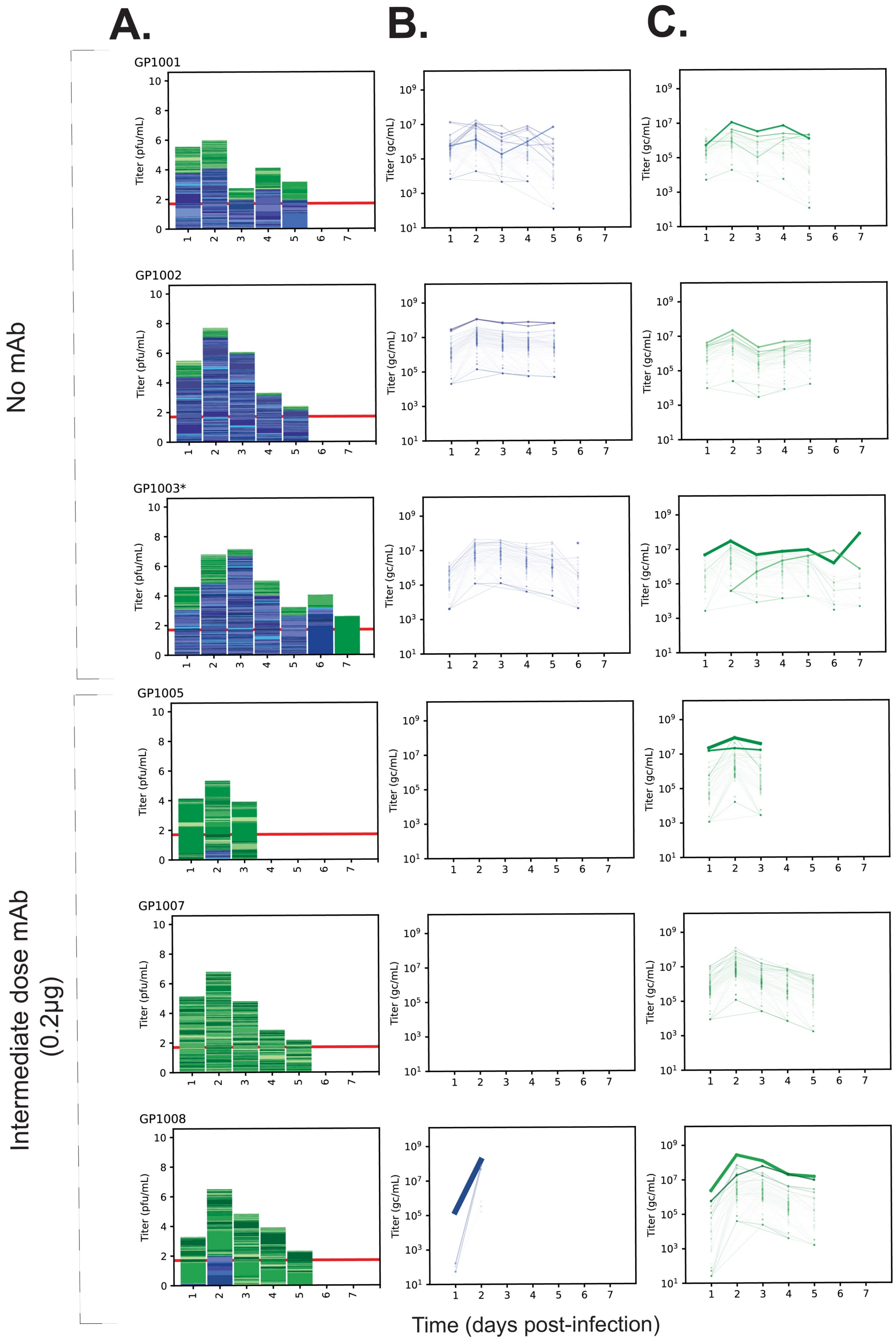

A.
B.
C.
No mAb
Intermediate dose mAb (0.2μg)
GP1001
GP1002
GP1003*
GP1005
GP1007
GP1008
Titer (pfu/mL)
Titer (gc/mL)
Time (days post-infection)

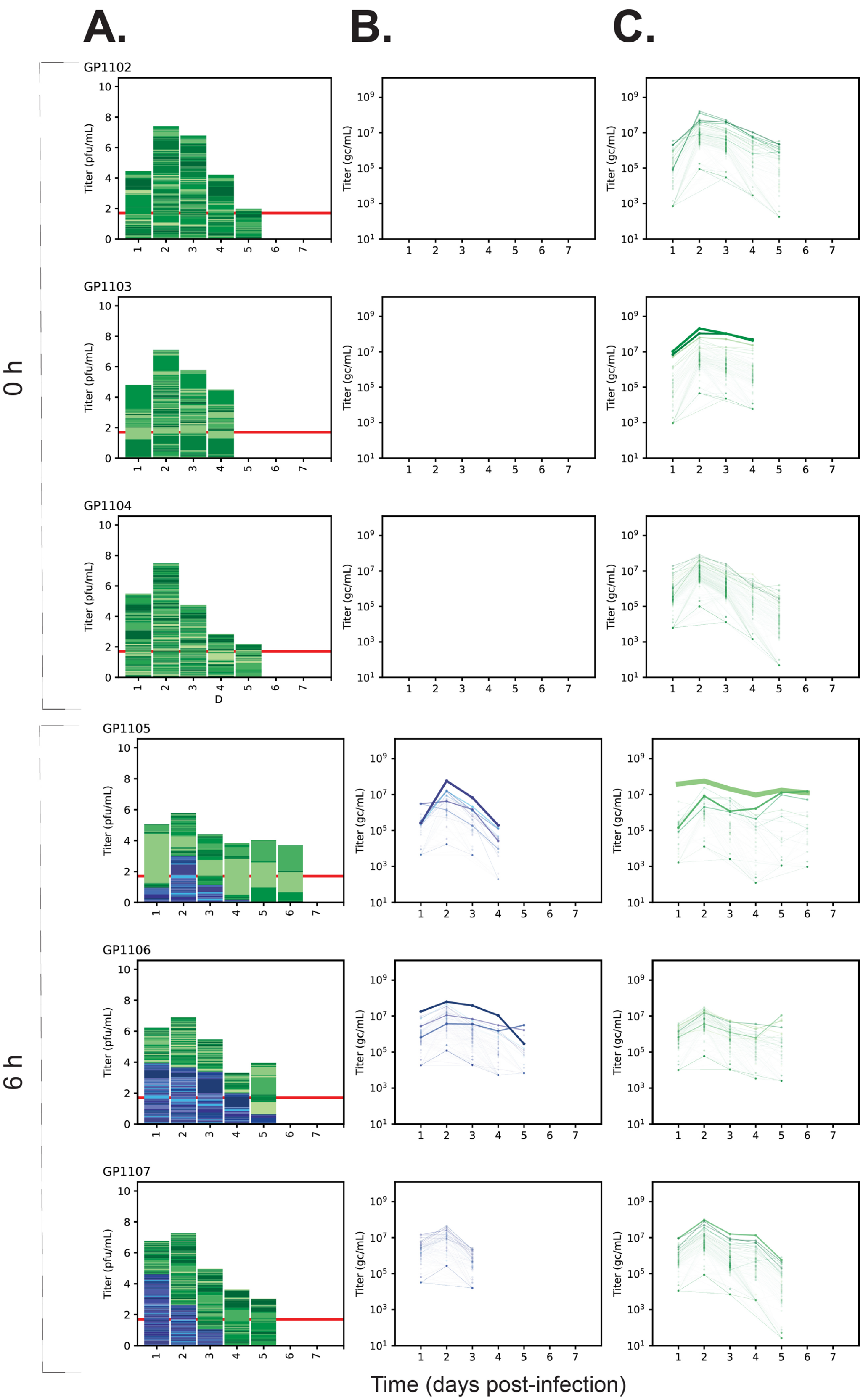
A.
B.
C.
0 h
6 h
GP1102
GP1103
GP1104
GP1105
GP1106
GP1107
Titer (pfu/mL)
Titer (gc/mL)
D
Time (days post-infection)

**Supplemental Figure 8: Antigenic selection is efficient when fitness costs are eased through the addition of second-site mutations. (Related to Figure 5).** Representative animals from these experimental conditions are shown in Figure 5 and remaining animals are shown here. Tx/12 HA WT BC1 virus is represented by blue/purple colors and Tx/12 HA A138S/F159S/N225D BC2 is represented by green colors. Animals were inoculated with 90% WT and 10% HA A138S/F159S/N225D BC viruses, and asterisk denotes animals in which *de novo* mutations arose and swept with a barcode. **(A)** Stacked bar plots where the total height of the column is scaled to viral titer ($\log_{10}$ pfu/ml). Each colored section within the column represents a unique barcode, and its height is the relative frequency within the sample. The total height of green bars and blue/purple bars is scaled to the gc/ml titer of each virus. Red horizontal line represents limit of detection of plaque assay (50 pfu/ml). **(B)** and **(C)** show Tx/12 HA WT and A138S/F159S/N225D individual barcodes over time, respectively. Each line is a unique barcode with its frequency scaled to the gc/ml titer. Line thickness is scaled to the maximum detected frequency of the barcode. Gaps denote results below the limit of detection. Dashed lines connect data separated by single days below the limit of detection.

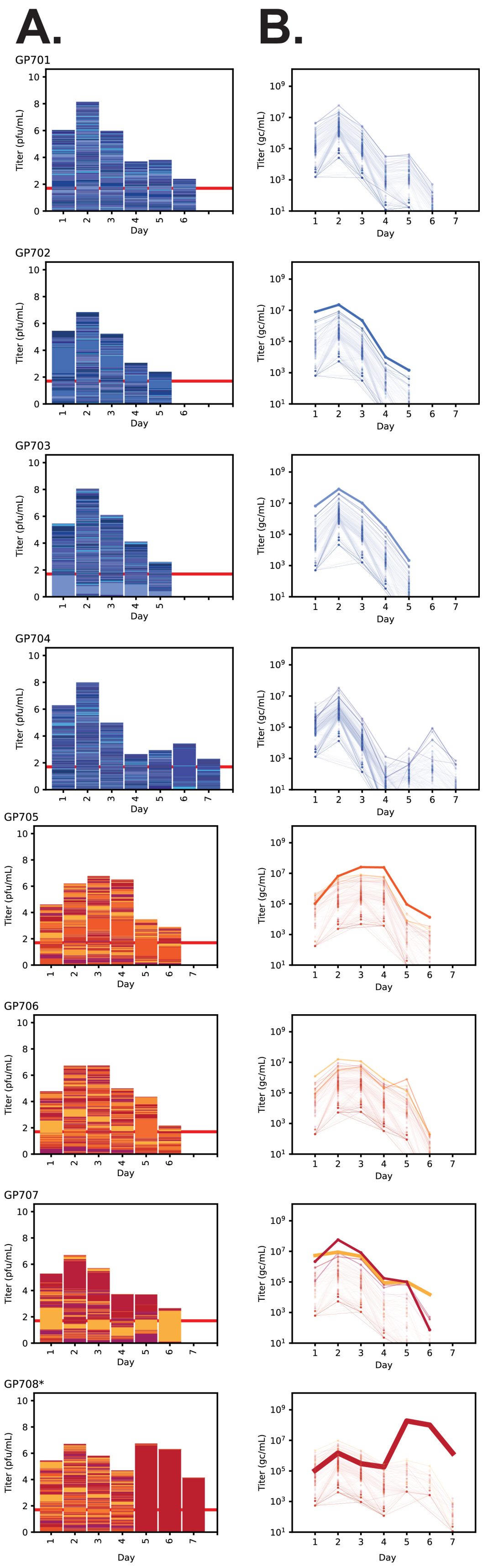
A.
B.
GP701
GP702
GP703
GP704
GP705
GP706
GP707
GP708*
Titer (pfu/mL)
Titer (gc/mL)
Day

**Supplemental Figure 9: Many barcodes are maintained over the course of infection in the absence of competition between alleles.** Data were obtained from animals mono-infected with Tx/12 WT or HA F159S BC viruses. **(A)** Stacked bar plots where the total height of the column is scaled to viral titer ($log_{10}$ pfu/ml). Each colored section within the column represents a unique barcode, and its height is the relative frequency within the sample. The total height of red/yellow bars and blue/purple bars is scaled to the gc/ml titer of each virus. Red horizontal line represents limit of detection of plaque assay (50 pfu/ml). **(B)** Tx/12 HA WT and F159S individual barcodes over time. Each line is a unique barcode with its frequency scaled to the gc/ml titer. Line thickness is scaled to the maximum detected frequency of the barcode. Gaps denote results below the limit of detection. Dashed lines connect data separated by single days that fell below the limit of detection.
