## Supplement 2 for "Within-host antigenic selection of influenza A virus dominates over stochasticity but is limited by fitness tradeoffs and timing of the immune response"

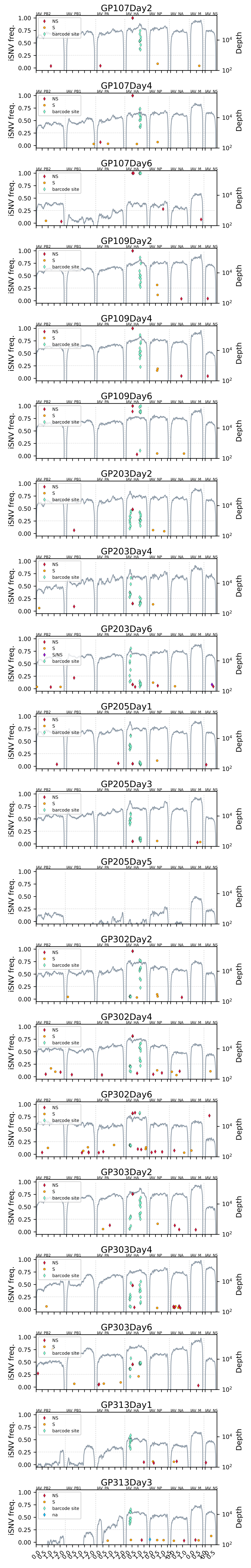
GP107Day2
GP107Day4
GP107Day6
GP109Day2
GP109Day4
GP109Day6
GP203Day2
GP203Day4
GP203Day6
GP205Day1
GP205Day3
GP205Day5
GP302Day2
GP302Day4
GP302Day6
GP303Day2
GP303Day4
GP303Day6
GP313Day1
GP313Day3
NS
S
S/NS
na
barcode site
iSNV freq.
Depth
IAV PB2
IAV PB1
IAV PA
IAV HA
IAV NP
IAV NA
IAV M
IAV NS

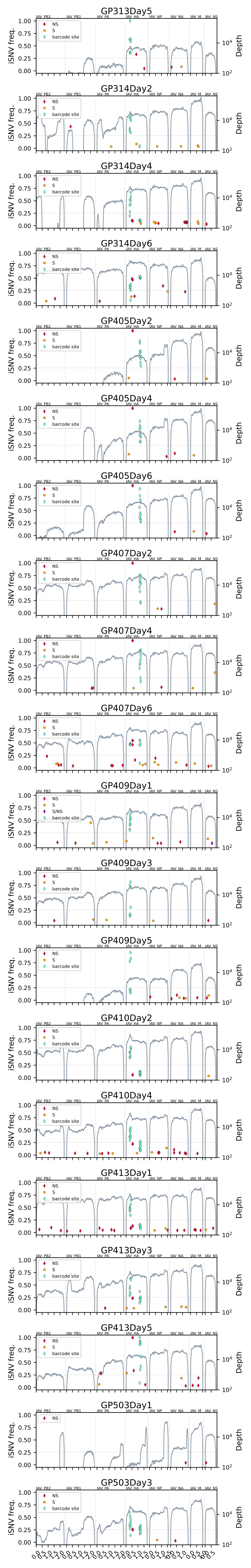
GP313Day5
GP314Day2
GP314Day4
GP314Day6
GP405Day2
GP405Day4
GP405Day6
GP407Day2
GP407Day4
GP407Day6
GP409Day1
GP409Day3
GP409Day5
GP410Day2
GP410Day4
GP413Day1
GP413Day3
GP413Day5
GP503Day1
GP503Day3
iSNV freq.
Depth
NS
S
S/NS
barcode site
IAV_PB2
IAV_PB1
IAV_PA
IAV_HA
IAV_NP
IAV_NA
IAV_M
IAV_NS

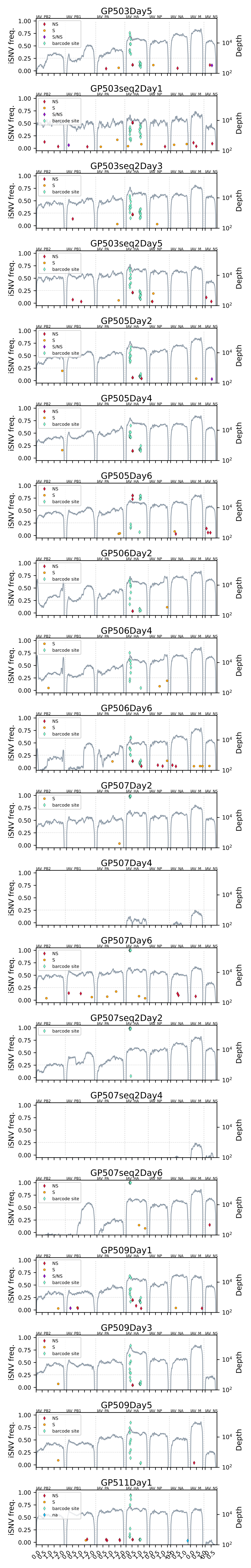

GP503Day5
GP503seq2Day1
GP503seq2Day3
GP503seq2Day5
GP505Day2
GP505Day4
GP505Day6
GP506Day2
GP506Day4
GP506Day6
GP507Day2
GP507Day4
GP507Day6
GP507seq2Day2
GP507seq2Day4
GP507seq2Day6
GP509Day1
GP509Day3
GP509Day5
GP511Day1
iSNV freq.
Depth
IAV_PB2
IAV_PB1
IAV_PA
IAV_HA
IAV_NP
IAV_NA
IAV_M
IAV_NS
NS
S
S/NS
barcode site
na

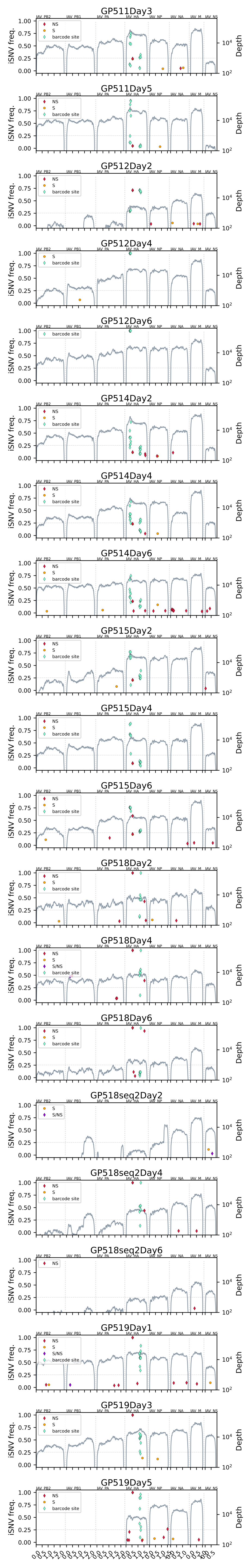

GP511Day3
GP511Day5
GP512Day2
GP512Day4
GP512Day6
GP514Day2
GP514Day4
GP514Day6
GP515Day2
GP515Day4
GP515Day6
GP518Day2
GP518Day4
GP518Day6
GP518seq2Day2
GP518seq2Day4
GP518seq2Day6
GP519Day1
GP519Day3
GP519Day5
iSNV freq.
Depth
NS
S
S/NS
barcode site
IAV_PB2
IAV_PB1
IAV_PA
IAV_HA
IAV_NP
IAV_NA
IAV_M
IAV_NS

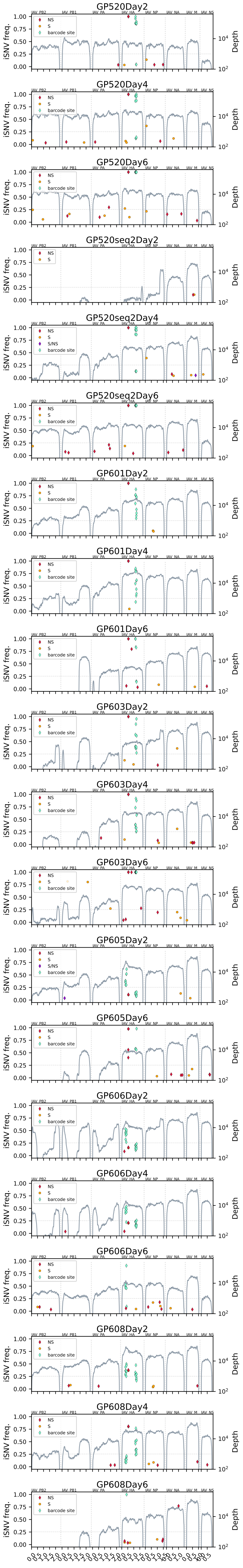
GP520Day2
GP520Day4
GP520Day6
GP520seq2Day2
GP520seq2Day4
GP520seq2Day6
GP601Day2
GP601Day4
GP601Day6
GP603Day2
GP603Day4
GP603Day6
GP605Day2
GP605Day6
GP606Day2
GP606Day4
GP606Day6
GP608Day2
GP608Day4
GP608Day6
iSNV freq.
Depth
NS
S
S/NS
barcode site
IAV_PB2
IAV_PB1
IAV_PA
IAV_HA
IAV_NP
IAV_NA
IAV_M
IAV_NS

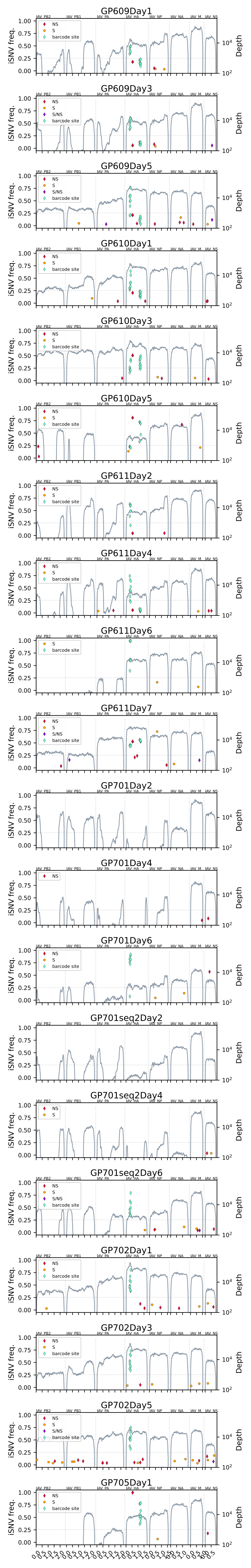
GP609Day1
GP609Day3
GP609Day5
GP610Day1
GP610Day3
GP610Day5
GP611Day2
GP611Day4
GP611Day6
GP611Day7
GP701Day2
GP701Day4
GP701Day6
GP701seq2Day2
GP701seq2Day4
GP701seq2Day6
GP702Day1
GP702Day3
GP702Day5
GP705Day1
iSNV freq.
Depth
NS
S
S/NS
barcode site
IAV_PB2
IAV_PB1
IAV_PA
IAV_HA
IAV_NP
IAV_NA
IAV_M
IAV_NS

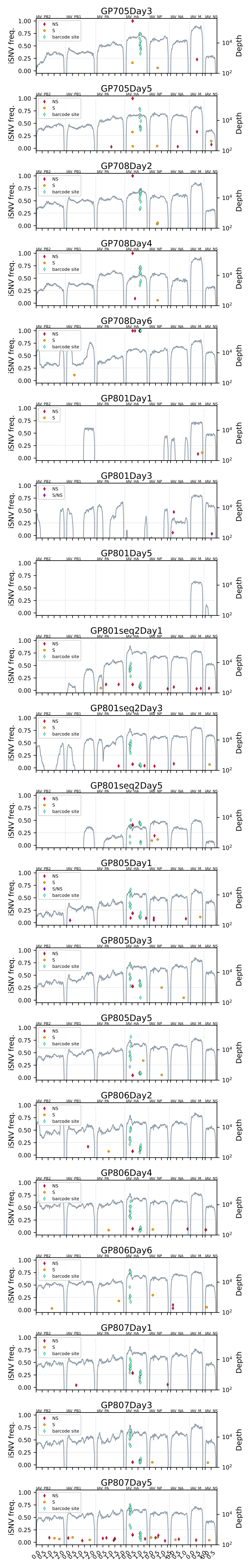

GP705Day3
GP705Day5
GP708Day2
GP708Day4
GP708Day6
GP801Day1
GP801Day3
GP801Day5
GP801seq2Day1
GP801seq2Day3
GP801seq2Day5
GP805Day1
GP805Day3
GP805Day5
GP806Day2
GP806Day4
GP806Day6
GP807Day1
GP807Day3
GP807Day5
IAV_PB2
IAV_PB1
IAV_PA
IAV_HA
IAV_NP
IAV_NA
IAV_M
IAV_NS
NS
S
S/NS
barcode site
iSNV freq.
Depth

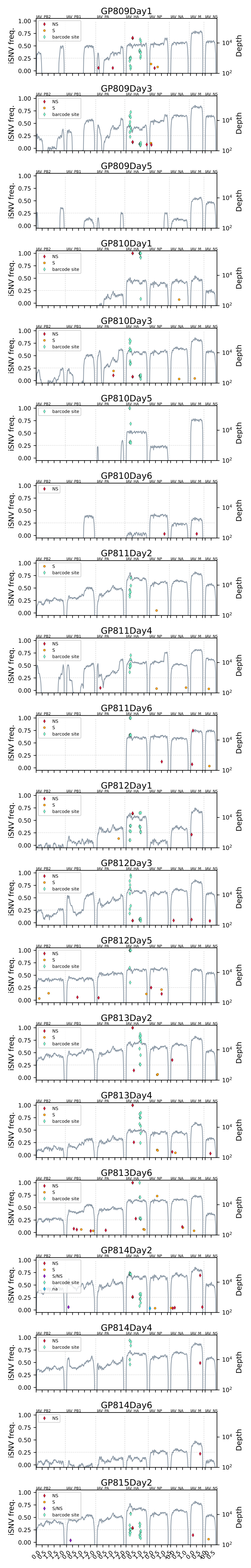

GP809Day1
GP809Day3
GP809Day5
GP810Day1
GP810Day3
GP810Day5
GP810Day6
GP811Day2
GP811Day4
GP811Day6
GP812Day1
GP812Day3
GP812Day5
GP813Day2
GP813Day4
GP813Day6
GP814Day2
GP814Day4
GP814Day6
GP815Day2
iSNV freq.
Depth
NS
S
S/NS
barcode site
na
IAV PB2
IAV PB1
IAV PA
IAV HA
IAV NP
IAV NA
IAV M
IAV NS

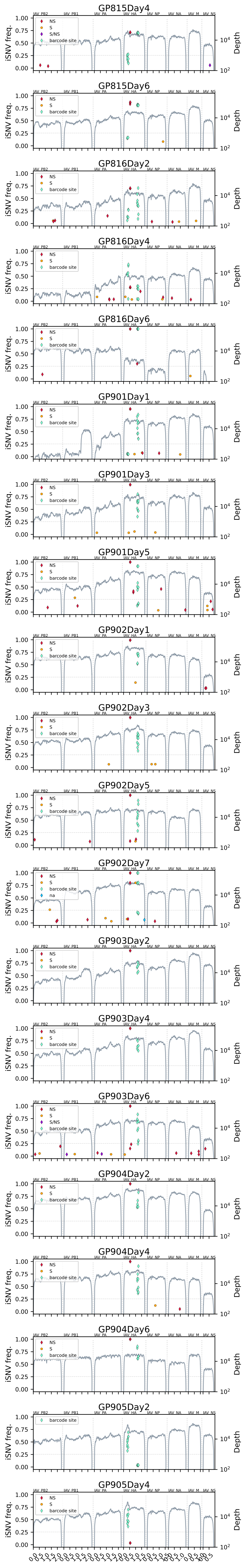
GP815Day4
GP815Day6
GP816Day2
GP816Day4
GP816Day6
GP901Day1
GP901Day3
GP901Day5
GP902Day1
GP902Day3
GP902Day5
GP902Day7
GP903Day2
GP903Day4
GP903Day6
GP904Day2
GP904Day4
GP904Day6
GP905Day2
GP905Day4
iSNV freq.
Depth
NS
S
S/NS
barcode site
na
IAV PB2
IAV PB1
IAV PA
IAV HA
IAV NP
IAV NA
IAV M
IAV NS

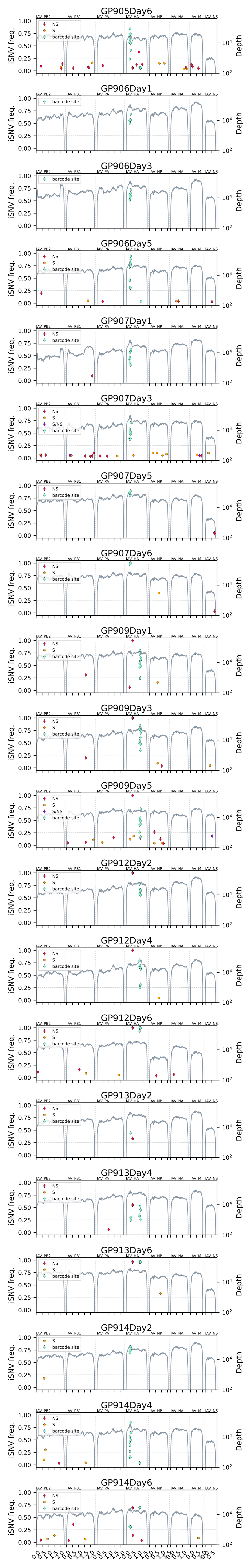
GP905Day6
GP906Day1
GP906Day3
GP906Day5
GP907Day1
GP907Day3
GP907Day5
GP907Day6
GP909Day1
GP909Day3
GP909Day5
GP912Day2
GP912Day4
GP912Day6
GP913Day2
GP913Day4
GP913Day6
GP914Day2
GP914Day4
GP914Day6
IAV PB2
IAV PB1
IAV PA
IAV HA
IAV NP
IAV NA
IAV M
IAV NS
NS
S
S/NS
barcode site
iSNV freq.
Depth

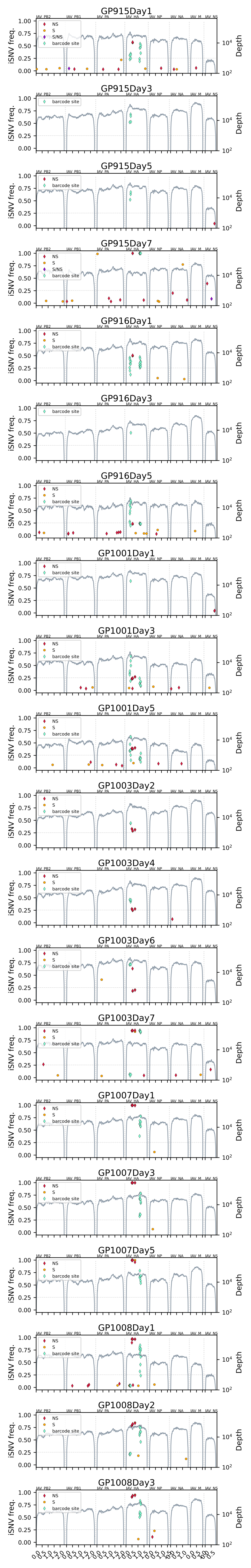
GP915Day1
GP915Day3
GP915Day5
GP915Day7
GP916Day1
GP916Day3
GP916Day5
GP1001Day1
GP1001Day3
GP1001Day5
GP1003Day2
GP1003Day4
GP1003Day6
GP1003Day7
GP1007Day1
GP1007Day3
GP1007Day5
GP1008Day1
GP1008Day2
GP1008Day3
iSNV freq.
Depth
NS
S
S/NS
barcode site
IAV PB2
IAV PB1
IAV PA
IAV HA
IAV NP
IAV NA
IAV M
IAV NS

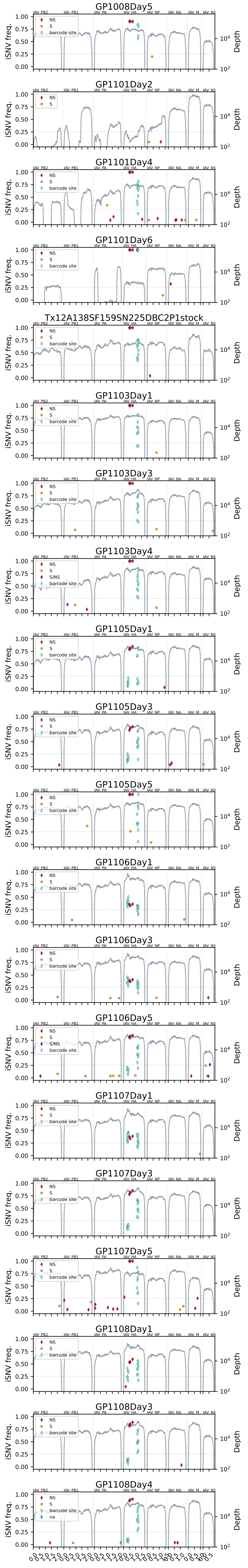

GP1008Day5
GP1101Day2
GP1101Day4
GP1101Day6
Tx12A138SF159SN225DBC2P1stock
GP1103Day1
GP1103Day3
GP1103Day4
GP1105Day1
GP1105Day3
GP1105Day5
GP1106Day1
GP1106Day3
GP1106Day5
GP1107Day1
GP1107Day3
GP1107Day5
GP1108Day1
GP1108Day3
GP1108Day4
IAV PB2
IAV PB1
IAV PA
IAV HA
IAV NP
IAV NA
IAV M
IAV NS
NS
S
S/NS
barcode site
na
iSNV freq.
Depth

**Supplemental Figure 6: Whole genome sequencing reveals *de novo* mutations that arise and increase in frequency during infection in a subset of animals.** Whole genome sequencing of guinea pig nasal wash samples. Categorized iSNVs (S=synonymous, NS=nonsynonymous, na=UTR) are plotted at their frequency on the left axis and sequencing coverage (gray line) is plotted on the right axis. Segments are identified above each trace.
