## Supplementary material for "Within-host antigenic selection of influenza A virus dominates over stochasticity but is limited by fitness tradeoffs and timing of the immune response": Table S3

**Table S3. Barcode mutations, related to Figure 1A**

| Barcode | Nucleotide position | Amino acid position | Codon | Amino acid |
|---|---|---|---|---|
| HA BC1 | 308 | 93 | GAT/GAC | D |
| HA BC1 | 314 | 95 | TTC/TTT | F |
| HA BC1 | 320 | 97 | AAT/AAC | N |
| HA BC1 | 326 | 99 | AAA/AAG | K |
| HA BC1 | 338 | 103 | TTT/TTC | F |
| HA BC1 | 347 | 106 | CGA/CGC | R |
| HA BC1 | 377 | 116 | TAT/TAC | Y |
| HA BC1 | 386 | 119 | CCG/CCA | P |
| HA BC1 | 395 | 122 | GCC/GCA | A |
| HA BC1 | 404 | 125 | AGG/AGA | R |
| HA BC2 | 1133 | 368 | GGT/GGC | G |
| HA BC2 | 1139 | 370 | AGG/AGA | R |
| HA BC2 | 1154 | 375 | GAG/GAA | E |
| HA BC2 | 1160 | 377 | AGA/AGG | R |
| HA BC2 | 1178 | 383 | CTT/CTC | L |
| HA BC2 | 1202 | 391 | GAT/GAC | D |
| HA BC2 | 1214 | 395 | GGG/GGA | G |
| HA BC2 | 1224 | 399 | CGA/AGA | R |
| HA BC2 | 1229 | 400 | TTG/TTA | L |
| HA BC2 | 1235 | 402 | GGG/GGA | G |
