## Supplementary material for "Within-host antigenic selection of influenza A virus dominates over stochasticity but is limited by fitness tradeoffs and timing of the immune response": Table S4

**Table S4. Oligonucleotides list**

| |
|---|
| Primer: Uni12/Inf-1 GGGGGGAGCAAAAGCAGG |
| Primer: Uni12/Inf-3 GGGGGGAGCGAAAGCAGG |
| Primer: Uni13/Inf-1 CGGGTTATTAGTAGAAACAAGG |
| Primer: Tx12_ddPCR_primerF CCGGCACACTGGAGTTTAACAATG |
| Primer: Tx12_ddPCR_primerR GGCATAGTCACGTTCAATGCTGG |
| Probe: Tx12_ddPCR_WT_probe TATTT+GAAGTT+TAAGT+G+G+GT+CAA |
| Probe: Tx12_ddPCR_HAF159S_probe TATTTA+CTGTTTAA+GT+G+G+GT+CAA |
| Primer: Tx12_HABC1seq_F<br>TCGTCGGCAGCGTCAGATGTGTATAAGAGACAGCTATTGGGAGACCCTCAGTGT |
| Primer: Tx12_HABC1seq_R<br>GTCTCGTGGGCTCGGAGATGTGTATAAGAGACAGGATGAGGCAACTAGTGA |
| Primer: Tx12_HABC2seq_F<br>TCGTCGGCAGCGTCAGATGTGTATAAGAGACAGGAATGGTGGATGGTTGGTAC |
| Primer: Tx12_HABC2seq_R<br>GTCTCGTGGGCTCGGAGATGTGTATAAGAGACAGGATGGAATTTCTCGTTGGTTTT |
| Primer: Tx12_HABC1_400down CACTAGTTGCCTCATCCGGCACACT |
| Primer: Tx12_HABC1_298up CACTGAGGGTCTCCCAATAGAGCAT |
| Primer: Tx12_HABC2_1234down CCAACGAGAAATTCCATCAGATTGA |
| Primer: Tx12_HABC2_1123up TACCAACCATCCACCATTCCCTCCC |
| Primer: Tx12_HABC1_78F CTTCCTGGAAATGACAATAGCACGGC |
| Primer: Tx12_HABC2_1019F GGCAACAGGAATGCGGAATGTACC |
| Ultramer: Tx12_HABC1_ultramer_F<br>AGATGCTCTATTGGGAGACCCTCAGTGTGAYGGCTTYCAAAAYAAGAARTGGGACCTTTTYGTTGAACGMAGCAAAGCCTACAGCAACTGTTACCCTTAYGATGTGCCRGATTATGCMTCCCTTAGRTCACTAGTTGCCTCATCCGGCACACTGGA |

| |
|---|
| Ultramer: Tx12_HABC1_ultramer_R<br>TCCAGTGTGCCGGATGAGGCAACTAGTGAYCTAAGGGAKGCATAATCYGGCACA<br>TCRTAAGGGTAACAGTTGCTGTAGGCTTTGCTKCGTTCAACRAAAAGGTCCCAYT<br>TCTTRTTTTGRAAGCCRTCACACTGAGGGTCTCCCAATAGAGCATCT |
| Ultramer: Tx12_HABC2_ultramer_F<br>TTGGGAGGGAATGGTGGATGGTTGGTACGGYTTCAGRCATCAAAATTCTGARGG<br>AAGRGGACAAGCAGCAGATCTYAAAAGCACTCAAGCAGCAATCGAYCAAATCAA<br>TGGRAAGCTGAATMGATTRATCGGRAAAACCAACGAGAAATTCCATCAGATTGA |
| Ultramer: Tx12_HABC2_ultramer_R<br>TCAATCTGATGGAATTTCTCGTTGGTTTTYCCGATYAATCKATTCAGCTTYCCATT<br>GATTTGRTCGATTGCTGCTTGAGTGCTTTTRAGATCTGCTGCTTGTCCYCTTCCY<br>TCAGAATTTTGATGYCTGAARCCGTACCAACCATCCACCATTCCCTCCCAA |
| Primer: UnivF(A)+6 GCGCGCAGCAAAAGCAGG |
| Primer: UnivF(G)+6 GCGCGCAGCGAAAGCAGG |
| Primer: Tx12_HABC1_XhoIF<br>CAGTGTGATGGCTTCCAAAATAAGAAACTCGAGCTTTTTGTTGAACG |
| Primer: Tx12_HABC1_XhoIR<br>CGTTCAACAAAAAGCTCGAGTTTCTTATTTTGGAAGCCATCACACTG |
| Primer: Tx12_HABC1_stop1F GTTGAACGAAGCAAATAGTACAGCAACTGTTAC |
| Primer: Tx12_HABC1_stop1R GTAACAGTTGCTGTACTATTTGCTTCGTTCAAC |
| Primer: Tx12_HABC1_stop2F GCCGGATTATGCCTCCTAGAGGTCACTAGTTGC |
| Primer: Tx12_HABC1_stop2R GCAACTAGTGACCTCTAGGAGGCATAATCCGGC |
| Primer: Tx12_HABC2_XhoIF<br>AGGGAAGAGGACAAGCTCGAGATCTCAAAAGCACTC |

| |
|---|
| Primer: Tx12_HABC2_XhoIR GAGTGCTTTTGAGATCTCGAGCTTGTCCTCTTCCCT |
| Primer: Tx12_HABC2_stop1F GGTTTCAGGCATCAATAGTCTGAGGGAAGAGGA |
| Primer: Tx12_HABC2_stop1R TCCTCTTCCCTCAGACTATTGATGCCTGAAACC |
| Primer: Tx12_HABC2_stop2F CTCAAAAGCACTCAATGAGCAATCGATC |
| Primer: Tx12_HABC2_stop2R GATCGATTGCTCATTGAGTGCTTTTGAG |
